# α5β1 and αvβ3 integrins employ two distinct adhesion strengthening modes to respond to fibronectin stiffness within seconds of initiating adhesion

**DOI:** 10.64898/2026.07.09.737593

**Authors:** Nico Strohmeyer, Upnishad Sharma, Michele M. Nava, Gotthold Fläschner, Javier Casares Arias, Daniel J. Müller

## Abstract

The complex interplay between extracellular matrix stiffness and actomyosin contractility regulates long-term integrin-based adhesion and signaling in mammalian cells. However, how cells sense and respond to the stiffness of the environment during the first minutes of initiating adhesion remains elusive. Here, we show that fibroblasts upon initiating adhesion to fibronectin switch between two distinct mechanosensitive adhesion modes. The first mode of “slow” adhesion strengthening, shared between α5β1 and αvβ3 integrins and depends modestly on fibronectin stiffness. Fibroblasts adapt this slow adhesion strengthening mode, which is independent of actomyosin contractility and intracellular signaling, on soft fibronectin substrates (<5 kPa). On stiff fibronectin substrates (>5 kPa), however, α5β1 integrins but not αvβ3 integrins switch to a “fast” adhesion strengthening mode that considerably strengthens adhesion within seconds. The switch to the fast mode depends on myosin II-mediated contractility and a mechanosensitive signaling hub that includes the α5β1 integrin-FN catch bond, paxillin, and focal adhesion kinase. The mechanistic findings highlight the similarities and differences of integrin-type specific adhesion strengthening and intracellular regulation, which depend on mechanotransduction in fibroblasts during adhesion initiation.

## Introduction

In mammals, cell adhesion undergoes dynamic adjustment in response to the biochemical composition and biophysical properties of the surrounding extracellular matrix (ECM)^[1,2]^. The active response of cells to the ECM is essential for a variety of physiological processes, such as development, morphogenesis, homeostasis, and tissue regeneration^[3–6]^ by regulating manifold cellular properties including adhesion, mechanics, differentiation, and metabolism^[7,8]^. The non-physiological ECM remodeling has severe pathological consequences. In fact, the abnormal biochemical and biophysical properties of the ECM are the underlying mechanisms of many pertinent diseases, especially solid and fibrotic malignant tumors that assemble stiff ECMs^[9,10]^.

Cells employ α/β heterodimeric, transmembrane receptors called integrins to adhere to ECM proteins, and to sense ECM properties biochemically and biophysically^[11,12]^. In mammals, 24 different integrins are formed by 18 α- and eight β-subunits. Several integrins are commonly co-expressed by the same cell. The Arg-Gly-Asp (RGD) motif is found in different ECM proteins, including fibronectin (FN) and represents the main binding site for α5β1 and αvβ3 integrin^[13]^. Additionally, α5β1 integrins bind to the synergy (PHSRN) site in FN, which is reported to be important for their firm ligand binding and mechanotransduction^[14–16]^. Integrin ligand binding is determined by reversible and thermodynamically induced conformation changes between a bent-closed conformation with low ligand binding affinity and an extended-open conformation in which integrins bind to their ligand with high affinity^[17–20]^.

Additionally, in the extended-open conformation integrins recruit the intracellular adaptor proteins, kindlin and talin, to the intracellular tail of the β subunit that mediate bidirectional force transduction between actin and the ECM ligand^[21–24]^. Specific integrins can respond to these forces by exhibiting catch-slip bonds, which strengthen under force (catch regime) up to a force threshold after which the bond weakens^[25,26]^. The transduction of force to talin by either actin retrograde flow or actomyosin contractility induces the actin-binding talin rod domain to unfold and the exposure of cryptic binding sites for vinculin, which in turn stabilizes talin in the unfolded state and strengthens the integrin-actin connection^[27–29]^. Furthermore, many other intracellular proteins involved in mediating integrin adhesion (adhesome proteins) form directionally asymmetric catch bonds or are mechanosensitive, where force-induced structural changes modulate the protein activity or exposure of cryptic binding sites to adhesome proteins^[30–33]^. Thereby, the force application to the adhesion site is indispensable to mature small nascent adhesions to large focal adhesions and to induce increased lifetimes of integrin-ECM bonds, postulated as molecular clutch^[34–37]^. Hence, the force stressing the adhesion site, which correlates with the ECM stiffness, determines the maturation state, size, and composition of the adhesion site, rendering integrins to sense the mechanical properties of the extracellular environment^[38]^.

Besides the immense advancement in understanding integrin mechanotransduction of optically resolvable adhesion sites, it remains elusive how very early stages of adhesion initiation are regulated by extracellular biomechanical cues and intracellular signaling. Furthermore, whether molecular clutch dynamics are integrin-type specific is unclear. Here, we employ atomic force microscopy (AFM)-based single-cell force spectroscopy (SCFS)^[39,40]^ to investigate how α5β1 and αvβ3 integrins regulate cell adhesion strengthening in response to soft, stiffer, and stiff FN substrates within the first seconds to minutes of adhesion initiation. We find that αvβ3 and α5β1 integrins employ different mechanisms and time scales to respond to FN stiffness.

## Results

### α5β1 but not αvβ3 integrins employ different adhesion initiation modes

To investigate how α5β1 and αVβ3 integrins strengthen adhesion on stiff FN substrates, we used AFM-based SCFS to quantify the adhesion force of genetically engineered fibroblast lines that express specific FN-binding integrins (**Figure S1, Supporting Information**). We brought single, rounded cantilever-bound pKO-β1 (expressing FN-binding α5β1 integrins) or pKO-αv fibroblasts (expressing FN-binding αvβ3 integrins)^[41]^ in contact with FN fragments (FNIII7-10)^[13]^ harboring the RGD and synergy site, coated onto glass surfaces, allowed them to interact with the substrate for contact times of 5 to 360 s and retracted the cantilever to quantify the force needed to detach the fibroblasts. pKO-β1 fibroblasts established higher adhesion force than pKO-αv fibroblasts at all contact times except for 20 s (**Figure S2a, Supporting Information**)^[13,42]^. The difference in adhesion force between pKO-β1 and pKO- αv fibroblasts increased with contact time and maximized in an 8-fold higher adhesion force at 360 s. Already at contact times >60 s pKO-β1 fibroblast established adhesion force over a wide range, which we divided into subpopulations of low (<15 nN), medium (15 – 30 nN), and high (>30 nN) adhesion force. With increasing contact time, the fibroblasts increasingly populated the medium and high adhesion forces (**Figure S2b, Supporting Information**). In contrast, pKO-αv fibroblasts showed only a single population of low adhesion force at contact times up to 240 s and sparsely populated the medium adhesion force (≈18 %) at 360 s (**Figure S2c, Supporting Information**)

Next, we quantified the adhesion strengthening rate of pKO-β1 and pKO-αv fibroblasts to FNIII7-10, which describes the increase in adhesion force over time. While the adhesion strengthening of whole population of pKO-β1 and pKO-αv fibroblast were best described by linear functions (**Figure S2d, Supporting Information**), we examined how each subpopulation of the adhesion force of pKO-β1 and pKO-αv fibroblasts strengthened adhesion. For statistical validity, we increased the number of fibroblasts characterized by SCFS at contact times >60 s (**Figure S2e, Supporting Information**). The low adhesion force population in pKO-β1 and pKO-αv fibroblasts was also best described by a linear function (**Figure 1a**). Surprisingly, the medium and high adhesion force subpopulations strengthened sigmoidally. Within contact times of 20 s and 120 s, both populations the adhesion force increased ≈9-fold for the medium and ≈22-fold for the high adhesion force population. Beyond this contact time, fibroblasts strengthened adhesion at modest rates.

**Figure 1.**
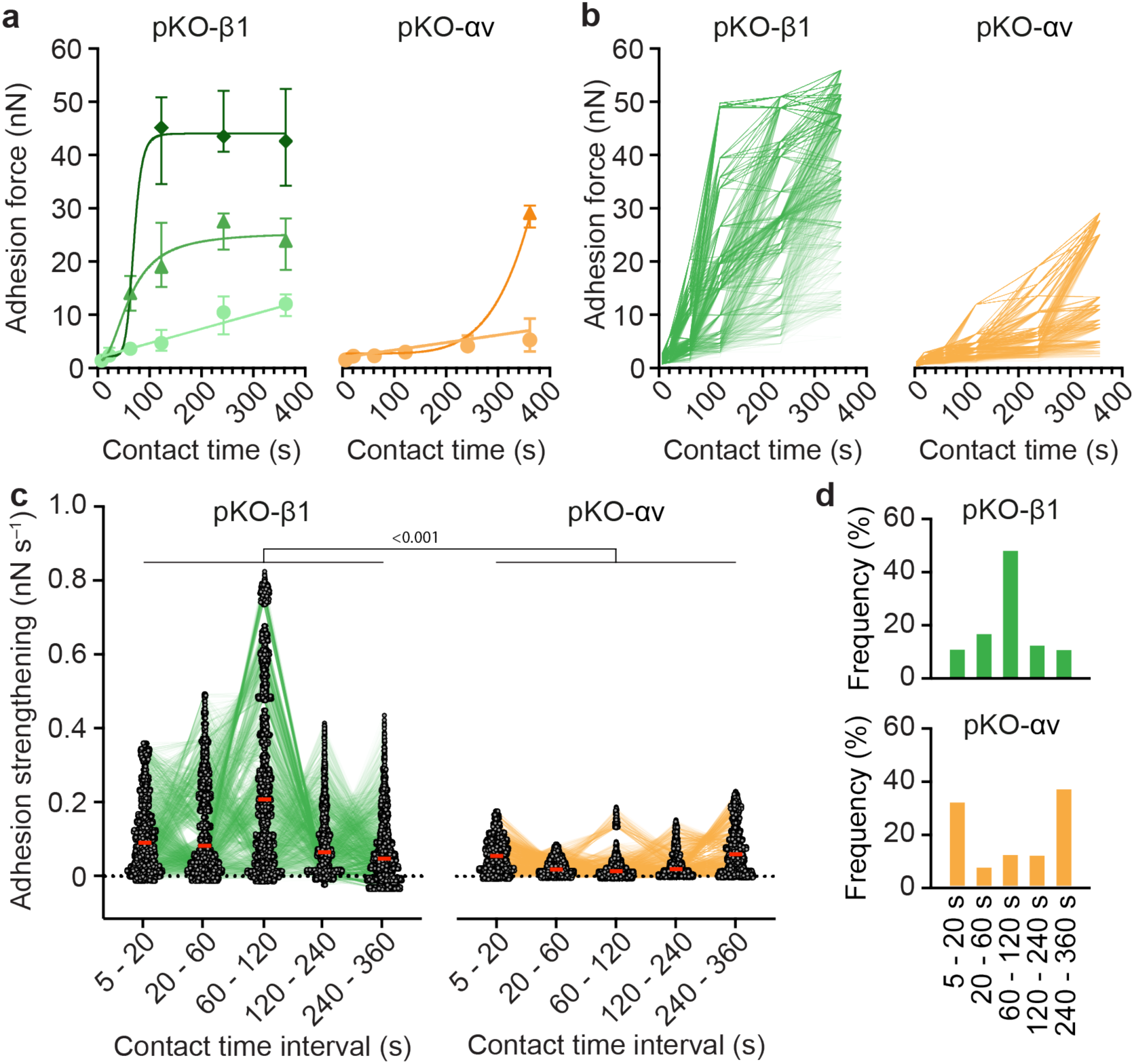
α5β1 but not αvβ3 integrins switch their adhesion strengthening mode to FNIII7-10 within the first minutes of initiating adhesion. **a)** Adhesion strengthening of (left) pKO-β1 and (right) pKO-αv fibroblast subpopulations to FNIII7-10 coated glass substrates. Adhesion forces of individual fibroblasts are shown in Figure S2e. Dots represent the median forces of the low, triangles of the medium, and rectangles of the high adhesion force subpopulations. Error bars depict the 95% confidence interval, and lines the fits. Low adhesion force populations are fitted best with linear functions, while the medium and low adhesion force populations are fitted best with sigmoidal functions. **b)** Simulated adhesion strengthening of individual (left) pKO-β1 and (right) pKO-αv fibroblasts to FNIII7-10 (*n*=4’000 each). Adhesion forces of data presented in Figure S2e were bootstrapped with the boundary condition of a maximally 10% adhesion force decrease to the previous contact time. **c)** Adhesion strengthening of modeled (left) pKO-β1 and (right) pKO-αv fibroblast to FNIII7-10 at given contact time intervals. Dots represent adhesion strengthening rates within a contact time interval, thin lines connect adhesion strengthening rates of individual fibroblasts, and red bars depict the median adhesion strengthening rate. P values comparing the adhesion strengthening rates of pKO-β1 and pKO-αv fibroblast at each contact time interval were calculated using two-sided Mann-Whitney tests. **d)** Fraction of simulated (top) pKO-β1 and (bottom) pKO-αv fibroblasts that show the highest adhesion strengthening rate at the given contact time interval during the first 360 s.

That fibroblasts populated low, medium and high adhesion force differently with increasing contact time and strengthened each adhesion force distribution differently indicates a complex behavior of adhesion strengthening. Hence we investigated the behavior of individual cells in more detail. Naturally, the fibroblasts attached to the cantilever spread over time. Thus, we could not apply SCFS to quantify the adhesion force of individual fibroblasts for all contact times. We hence simulated the adhesion strengthening of 4’000 single pKO-β1 or pKO-αv fibroblasts by bootstrapping to understand all possible adhesion strengthening trajectories from our SCFS experiments (**Figure 1b**; **Methods**). In the simulations, we allowed fibroblasts to establish maximally 10% lower adhesion force compared to the previous contact time. We implemented this boundary condition because SCFS experiments showed that fibroblasts unlikely reduce the adhesion force within the first 360 s of adhesion initiation (**Figure S2a, Supporting Information**).

The simulations showed that single pKO-β1 fibroblasts change the adhesion strengthening rate within the first 360 s of initiating adhesion. While ∼50% of pKO-β1 fibroblasts attained the maximum adhesion strengthening rate between 60 s and 120 s contact time the other ∼50% of pKO-β1 fibroblasts had their highest adhesion strengthening rate in the other contact time intervals. (**Figure 1c,d**). However, after reaching the maximum adhesion strengthening rate, individual pKO-β1 fibroblasts returned to modest adhesion strengthening rates. Compared to pKO-β1 fibroblasts, pKO-αv fibroblasts showed lower adhesion strengthening rates for all contact time intervals except for a slightly higher adhesion strengthening rate between 240 s and 360 s contact time, in which most pKO-αv fibroblasts showed their maximum adhesion strengthening rate (**Figure 1c,d**). Importantly, SCFS experiments of individual fibroblasts verified the different adhesion strengthening behaviors of pKO-β1 and pKO-αv fibroblasts (**Figure S2f, Supporting Information**).

In summary, simulations and experiments show that α5β1 integrins switch from a slow to a fast adhesion strengthening mode within the first 360 s of contact to FN, whereas αvβ3 integrins mainly remain in the low adhesion strengthening mode.

### Adhesion initiation regulation by talin-mediated integrin-actin engagement depends on integrin-type

To investigate how adhesion initiation of fibroblasts to FN is regulated by the actomyosin cortex, we characterized the adhesion strengthening of α5β1 or αvβ3 integrins that cannot engage the actin cytoskeleton *via* talin. To this end we quantified adhesion forces of talin1/2- depleted fibroblasts re-expressing the talin1 head domain (TKO+THD fibroblasts) to FNIII7- 10 in the presence of cilengitide to investigate α5β1 integrins^[42]^, or the β1-blocking antibody AIIB2 to investigate αv-class integrins^[43]^ (**Figure 2a**). Cilengitide-treated TKO+THD fibroblasts established lower adhesion forces at all contact times compared to pKO-β1 fibroblasts, and particularly at 120 s, 240 s and 360 s, for which the inability to engage α5β1 integrins to actin reduced the adhesion forces by a factor of ∼2, ∼5 and ∼7, respectively. Further, cilengitide-treated TKO+THD fibroblasts populated only the low adhesion force regimes and showed a linear adhesion strengthening (**Figure 2b** and **Figure S3a, Supporting Information**). The adhesion strengthening rate of cilengitide-treated TKO+THD fibroblasts to FNIII7-10 was lower compared to the low adhesion force population of pKO-β1 fibroblasts. AIIB2-treated TKO+THD fibroblasts established slightly higher adhesion forces to FNIII7-10 at 5 s contact time compared to pKO-αv fibroblasts (**Figure 2a**). At all other contact times, adhesion forces of AIIB2-treated TKO+THD and pKO-αv fibroblasts were similar and AIIB2-treated TKO+THD exclusively populated low adhesion forces (**Figure S3b, Supporting Information**). Hence, their adhesion strengthening was linear and indistinguishable from the low adhesion force population of pKO-αv fibroblasts (**Figure 2b**).

**Figure 2.**
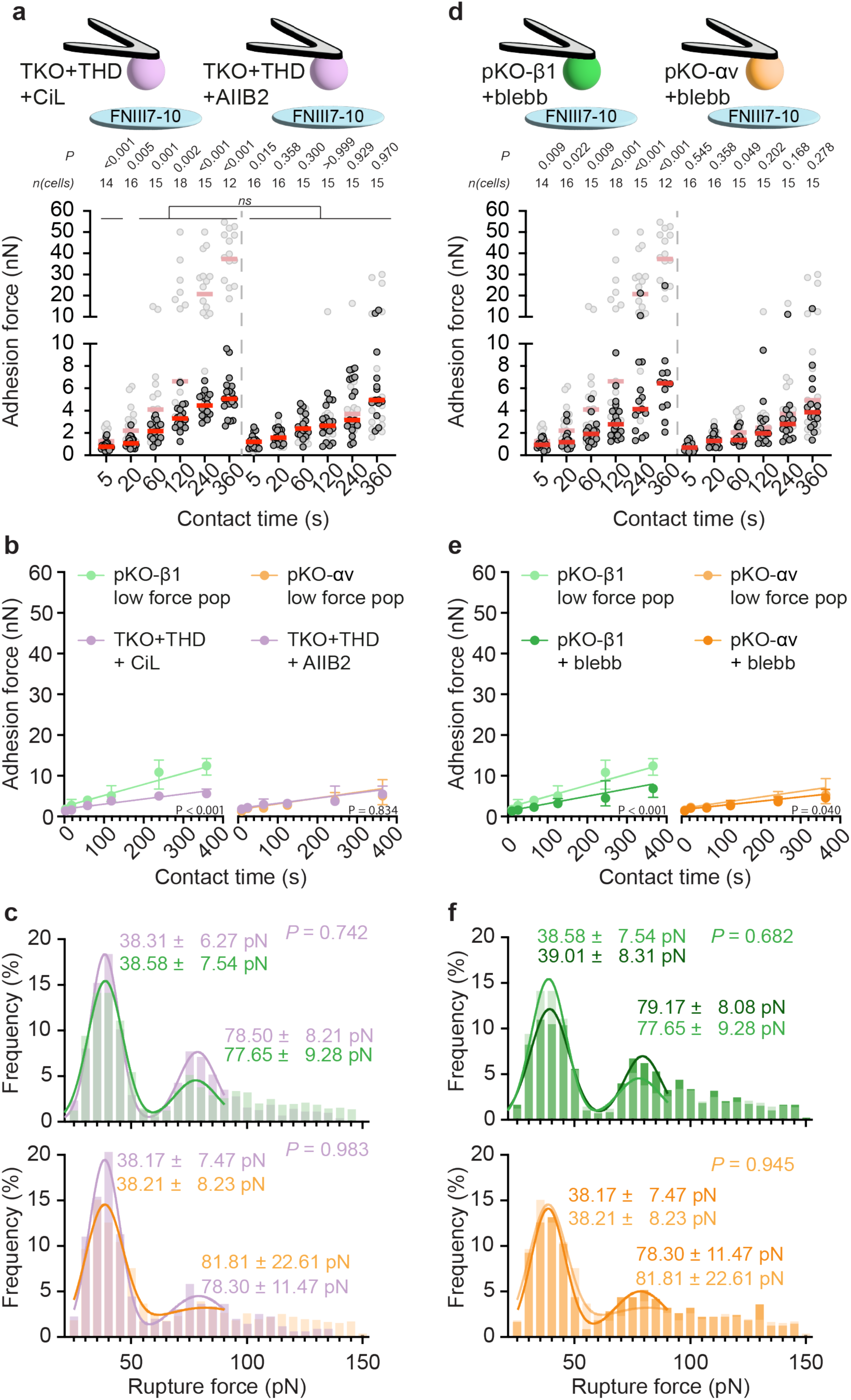
The actomyosin cortex regulates adhesion initiation of α5β1 but not αvβ3 integrins. **a**) Adhesion forces of talin1/2-depleted fibroblasts re-expressing the talin-head domain (TKO+THD fibroblasts) to FNIII7-10 in the presence of (left) cilengitide (CiL) and (right) AIIB2 at given contact times. Dots represent adhesion forces of individual fibroblasts, red bars the median, and n(cells) the number of fibroblasts tested. Adhesion forces of (left) pKO-β1 and (right) pKO-αv fibroblast are given in grey as reference (data taken from Figure S1a). P values comparing given and reference data were calculated using two-sided Mann-Whitney tests. **b)** Adhesion strengthening of TKO+THD fibroblasts to FNIII7-10 in the presence of (left) CiL and (right) AIIB2 was quantified as a slope of a linear function (lines) fitted to all adhesion forces at all contact times. Dots represent the mean adhesion force and bars the 95% confidence interval. The adhesion strengthening of (left) pKO-β1 fibroblasts or (right) pKO-αv fibroblasts are given as reference. The given *P* value compares the linear fit of both data sets and was calculated by extra sum-of-squares F-tests. **c)** Histograms (purple) with a bin size of 5 pN show the distribution of rupture forces of TKO+THD fibroblasts to FNIII7-10 in the presence of (top) CiL and (bottom) AIIB2 from FNIII7-10. For TKO+THD in presence of cilengitide *n* = 674 and for TKO+THD in presence of AIIB2 *n* = 367 single rupture events were analyzed. Lines show fits of a double Gaussian function to rupture force distributions between 20 and 90 pN. Rupture force distributions of untreated pKO-β1 fibroblasts (*n* = 1260 rupture events) and pKO-αv fibroblast (*n* = 1750 rupture events) are given in green and orange, respectively. P values compare means of the double Gaussian fits of given and reference data and were calculated by extra sum-of-squares F-tests. **d)** Adhesion forces of (left) pKO-β1 fibroblasts and (right) pKO-αv fibroblasts to FNIII7-10 in the presence of 20 µM blebbistatin (blebb) at given contact times. Data representation as in a). **e)** Adhesion strengthening rates of (left) pKO-β1 fibroblasts and (right) pKO-αv fibroblasts to FNIII7-10 in the presence of 20 µM blebbistatin. Data representation as in b). **f)** Rupture force distribution of blebbistatin- treated (left) pKO-β1 fibroblasts and (right) pKO-αv fibroblasts from FNIII7-10. For pKO-β1 fibroblasts *n* = 990 and for pKO-αv fibroblasts *n* = 515 single rupture events were analyzed.

Next, we investigated whether single integrin-ligand bonds are regulated by the engagement of integrins to the actomyosin cortex. Thereto, we analyzed force-distance (FD) curves recorded to quantify adhesion forces of pKO-β1, pKO-αv, AIIB2-treated TKO+THD and cilengitide-treated TKO+THD fibroblasts to FNIII7-10 for rupture forces occurring after the adhesion force peak (**Figure S1c, Supporting Information**). These rupture forces correspond to the maximum force single integrins can bear before detaching from the extracellular ligand^[39,40]^. The contact time-independence of rupture forces^[44]^ allowed us to pool FD curves recorded for all contact times for analysis. We only analyzed rupture forces ≤ 150 pN since higher rupture forces likely originate from the simultaneous unbinding of multiple integrins^[39]^. Additionally, we did not consider rupture forces < 25 pN since they typically fall into the noise level of our SCFS measurements. The rupture force distributions of pKO-β1 and pKO-αv fibroblasts showed multiple peaks and hence we fitted both distributions with a double Gaussian function (**Figure 2c**). The first rupture force peak was very similar among pKO-β1 and pKO-αv fibroblasts with means of 38.58 ± 7.54 pN and 38.23 ± 8.25 pN (mean ± SD), respectively. Both are in very good agreement with previously reported rupture force of single α5β1 and αvβ3 integrins^[15,39,44]^. The rupture force of cilengitide- and AIIB2 treated TKO+THD fibroblasts was similar to the rupture force of pKO-β1 and pKO-αv fibroblasts, respectively.

Together these results show that the switch of adhesion strengthening mode of α5β1 integrins depends on the integrin-actin engagement *via* talin, while the adhesion force established by αvβ3 integrins and the maximum force individual α5β1 and αvβ3 integrins can bear are independent of this actin engagement.

### Regulation of adhesion initiation by actomyosin contractility is integrin-type dependent

To understand how fibroblasts switch from the slow to the fast adhesion strengthening mode we inhibited myosinII-mediated actin contractility by blebbistatin in SCFS experiments with pKO-β1 or pKO-αv fibroblasts. MyosinII-inhibited pKO-β1 fibroblasts increasingly reduced the adhesion force to FNIII7-10 with contact time and most blebbistatin-treated pKO-β1 fibroblasts (>90%) populated low adhesion force regimes (**Figure 2d** and **Figure S3c, Supporting Information**). In addition, myosinII inhibition in pKO-β1 fibroblast considerably reduced the adhesion strengthening, which followed a linear function. Similar to cilengitide-treated TKO+THD fibroblasts, myosinII-inhibited pKO-β1 fibroblast also strengthened adhesion at a lower rate than untreated pKO-β1 fibroblast in the low adhesion force population (**Figure 2e**). In contrast, myosinII inhibition did not affect the adhesion force of pKO-αv fibroblasts to FNIII7- 10 (**Figure 2d**). However, myosinII-inhibited pKO-αv fibroblasts did not populate medium adhesion forces at contact times of 360 s (**Figure S3d, Supporting Information**). Further, myosinII inhibition in pKO-αv fibroblasts resulted in a slightly lower adhesion strengthening rate (**Figure 2e**). Lastly, the maximum force at which single αvβ3 or α5β1 integrins unbound from FNIII7-10 was independent of actomyosin contractility (**Figure 2f**).

### α5β1 integrin bridges intracellular contractility and FN synergy site to switch adhesion strengthening modes

Next, we investigated whether the synergy site in FN is required to switch from slow to fast adhesion strengthening mode of α5β1 integrins. To this end, we quantified the adhesion force of pKO-β1 fibroblasts to a FN fragment with a loss of function mutation in the synergy site (FNIII7-10mSyn)^[14,16]^. The inability of α5β1 integrins to bind to the synergy site drastically reduced the adhesion force of pKO-β1 fibroblasts (**Figure 3a**), which populated only low adhesion forces and their linear adhesion strengthening was similar to myosinII-inhibited pKO- β1 fibroblasts (**Figure S4a-c, Supporting Information**). Inhibiting myosinII-mediated contractility did not further reduce the adhesion force of pKO-β1 fibroblasts to FNIII7-10mSyn (**Figure 4a**). Importantly, pKO-β1 fibroblasts established higher adhesion force to FNIII7- 10mSyn than to a fibronectin fragment lacking the RGD domain (FNIII7-10ΔRGD; **Figure S4d, Supporting Information**)^[13,16]^, confirming their integrin specific binding to FNIII7-10mSyn. Furthermore, α5β1 integrins bound to FNIII7-10mSyn could bear ∼ 5 pN less before detaching from the ligand (**Figure 3b**), which corresponds to the loss of α5β1 integrin-FN catch bond formation^[14,15,25]^.

**Figure 3.**
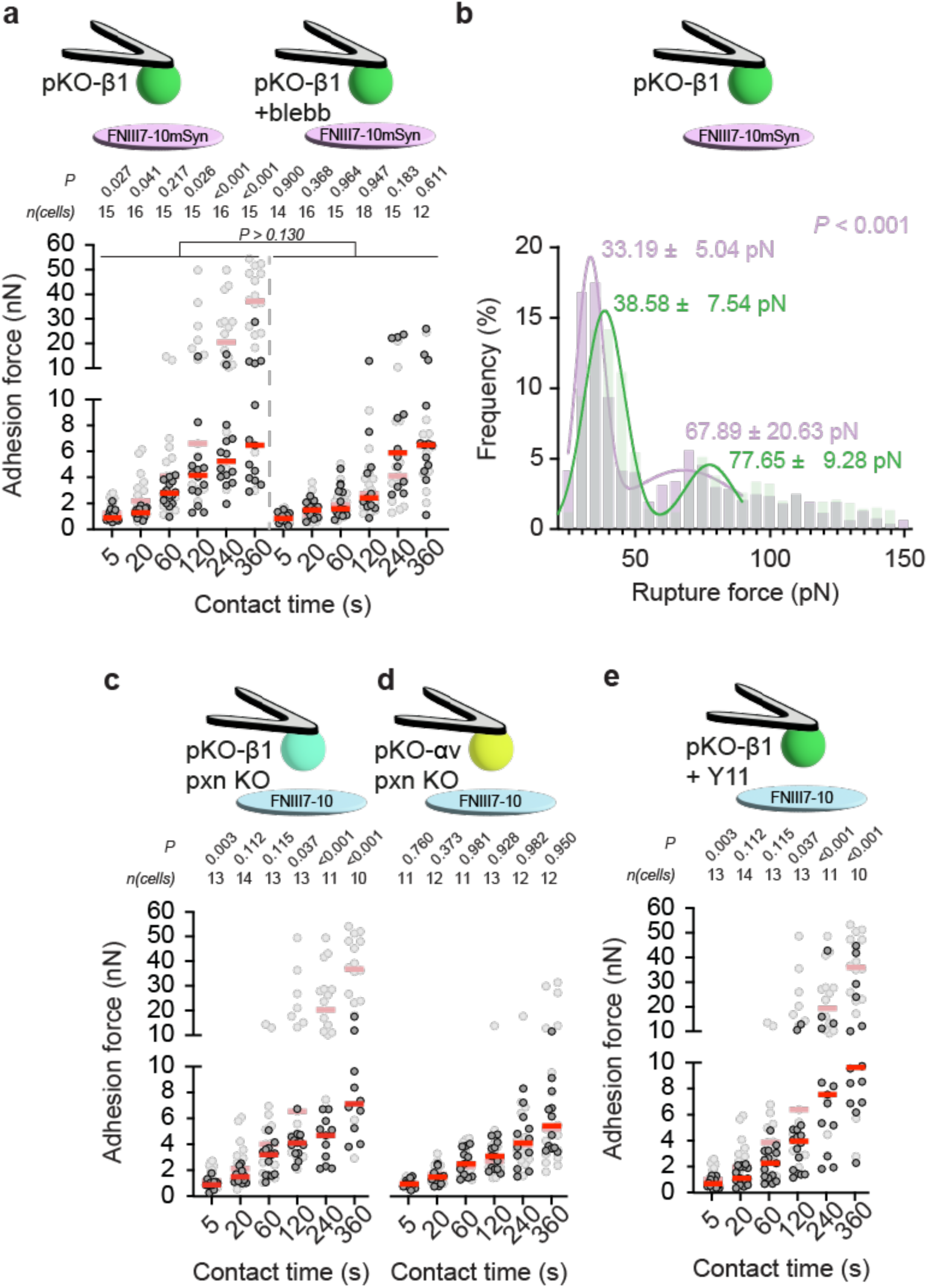
FN-α5β1 integrin catch bond, paxillin, and FAK signaling drive the adhesion strengthening mode switch in pKO-β1 fibroblasts. **a)** Adhesion forces of pKO-β1 fibroblasts to FNIII7-10mSyn in the (left) absence or (right) presence of 20 µM blebbistatin at given contact times. Dots represent adhesion forces of individual fibroblasts, red bars the median, and n(cells) the number of fibroblasts tested. Adhesion forces of pKO-β1 fibroblasts in the absence or presence of 20 µM blebbistatin to FNIII7-10 are given in grey as reference (data taken from Figure S1a). P values comparing given and reference data were calculated using two-sided Mann-Whitney tests. P values on bars compare adhesion forces of untreated and blebbistatin-treated pKO-β1 fibroblasts. **b)** Rupture forces distribution of pKO-β1 fibroblasts from FNIII7-10mSyn (*n* = 773, bin size 5 pN). Lines show fits of a double Gaussian function to rupture force distributions between 20 and 90 pN. Rupture force distributions of pKO-β1 fibroblast to FNIII7-10 is given in light green as reference (data taken from Figure 3c). Given P values compare means of the double gaussian fits of given and reference data and were calculated by extra sum-of-squares F-tests. **c-f)** Adhesion forces of c) pKO-β1 pxn KO fibroblasts, d) pKO-αv pxn KO fibroblasts, and e) pKO-β1 fibroblasts in the presence of 10 µM Y11 to FNIII7-10 at given contact time. Adhesion forces of pKO-β1 fibroblasts to FNIII7-10 are given in grey as reference (data taken from Figure 1). Data representation as described in a).

**Figure 4.**
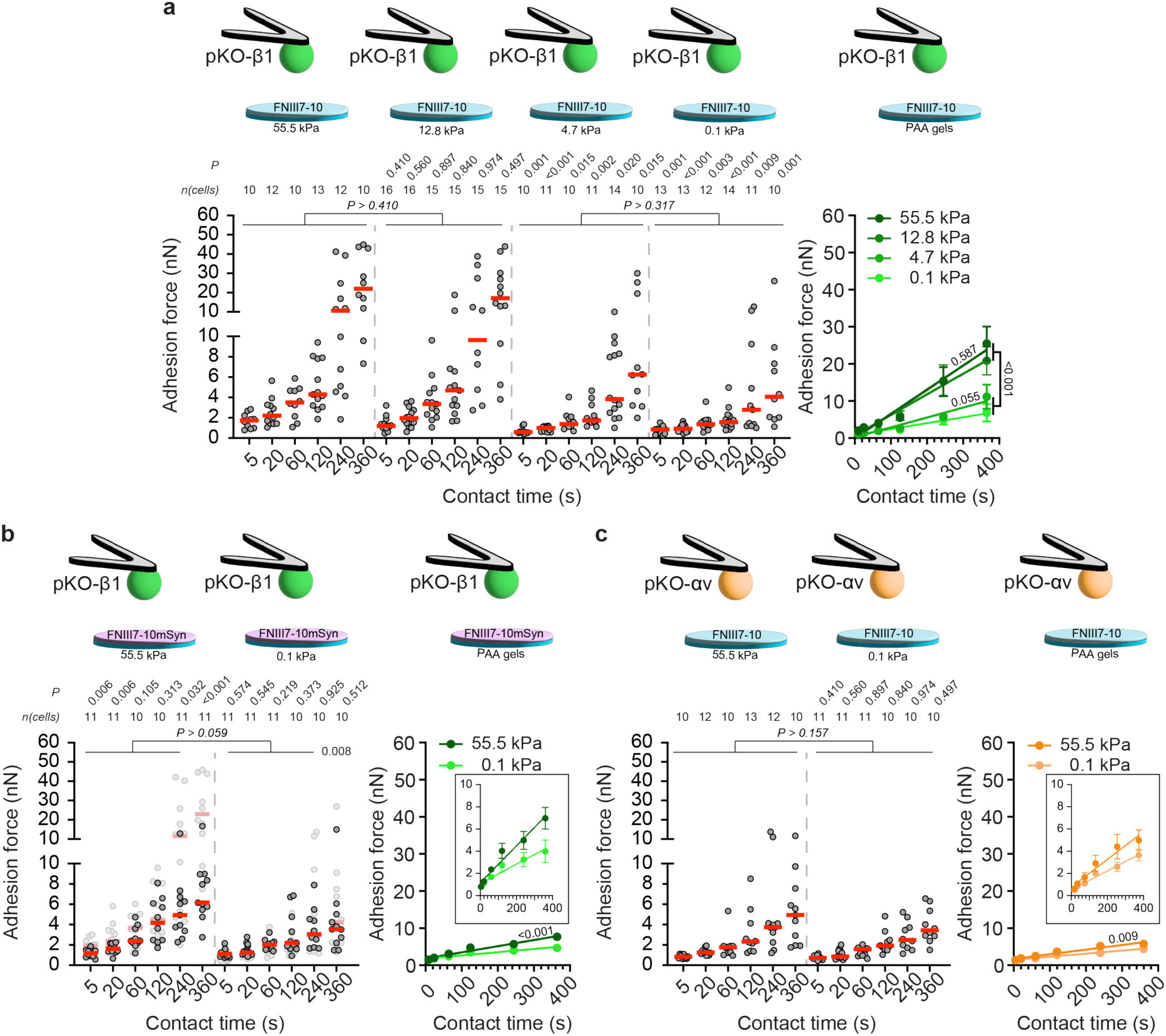
pKO-β1 and pKO-αv fibroblasts differently sense and respond to FN stiffness within seconds. Adhesion forces and strengthening of a) pKO-β1 fibroblasts to FNIII7-10 functionalized, b) pKO-β1 fibroblasts FNIII7-10mSyn-functionalized, or c) pKO-αv fibroblasts to FNIII7-10-functionalized PAA gels with given stiffness at given contact times. Dots represent adhesion forces of individual fibroblasts, red bars the medians, and *n*(cells) the number of fibroblasts tested. Reference data is given in the background in b) depicts adhesion forces of pKO-β1 fibroblasts FNIII7-10-functionalized PAA gels of the respective stiffness. P values in a) and d) compare given adhesion forces with a) pKO-β1 and c) pKO-αv fibroblasts to adhesion forces to the stiffest FNIII7-10-functionalized PAA gel or in b) given data with reference data. P values on lines compare adhesion forces of respective contact times of the indicated conditions. P values were calculated using two-sided Mann-Whitney tests. Adhesion strengthening was quantified by fitting a linear function to all adhesion forces for all contact time. Dots represent the mean adhesion forces, the error bars the standard error of the mean and the line depicts the linear fit. The given *P* values on the fits compare the linear fits and the P value on the lines compare the linear fits of the indicated conditions. P values were calculated by extra sum-of-squares F-tests.

Together the results show that the α5β1 integrin-FN catch bond and actomyosin contractility are part of the pathway that facilitates the fast adhesion strengthening mode of α5β1 integrins.

### Paxillin recruitment and focal adhesion kinase signaling regulate adhesion initiation in an integrin specific manner

To investigate molecular regulators of adhesion initiation, we targeted the integrin-type specific role of paxillin^[22]^ and the mechanosensitive focal adhesion kinase (FAK)^[14,16,45,46]^. We quantified the adhesion force of paxillin-null pKO-β1 (pKO-β1 pxn KO)^[47]^ and pKO-αv pxn KO fibroblasts as well as FAK-inhibited pKO-β1 fibroblasts to FNIII7-10 (**Figure 3c,d**). Depletion of paxillin in pKO-β1 fibroblasts reduced the adhesion force to FNIII7-10 at 5 s and ≥ 120 s contact time (**Figure 3c**). Importantly, pKO-β1 pxn KO fibroblasts populated only low adhesion forces and strengthened adhesion indistinguishable from myosinII-inhibited pKO-β1 fibroblasts (**Figure S5a-c, Supporting Information**). FAK-inhibition reduced the adhesion force of pKO-β1 fibroblasts to FNIII7-10 at all contact times (**Figure 3d**). However, at longer contact times, FAK-inhibited pKO-β1 fibroblasts established higher adhesion force to FNIII7- 10 than pKO-β1 pxn KO fibroblasts. Interestingly, while most FAK-inhibited pKO-β1 fibroblasts populated low adhesion forces, some fibroblasts populated medium and high adhesion forces (**Figure S5d, Supporting Information**).

### α5β1 and αvβ3 integrins employ different mechanisms to respond to FN stiffness during adhesion initiation

Finally, we investigated whether pKO-β1 or pKO-αv fibroblasts sense and respond to FN stiffness during adhesion initiation. As substrate we employed FNIII7-10 functionalized polyacrylamide (PAA) gels that cover a stiffness range of a variety of ECMs found in tissues^[48,49]^. Using the Hertz model^[50]^, we quantified the Young’s moduli of the PAA gels to be 0.13 ± 0.04 kPa (mean ± SD), 4.65 ± 1.00 kPa, 12.76 ± 2.03 kPa, and 55.45 ± 10.01 kPa for PAA gels (from here on called 0.1 kPa; 4.7 kPa; 12.8 kPa; 55.5 kPa gels, respectively; **Figure S6a, Supporting Information**). We adjusted the concentration of FNIII7-10 used to functionalize the differently stiff PAA gels to ensure similar ligand densities (**Methods**; **Figure S6b,c, Supporting Information**). pKO-β1 fibroblasts responded to the stiffness of PAA gels by increasing their spreading area 60 min after seeding (**Figure S6d**). This observation confirms that pKO-β1 fibroblasts respond to the stiffness of the FNIII7-10-functionalized PAA gels in long-term adhesion^[51]^.

We next quantified the adhesion force of pKO-β1 to ligand FNIII7-10 functionalized PAA gels (**Figure 4a**). Like for pKO-β1 fibroblasts adhering to FNIII7-10-coated glass, we observed subpopulations of low, medium, and high adhesion force (**Figure S7a, Supporting Information**). However, on FNIII7-10-functionalized PAA gels the medium and high adhesion force subpopulations only appeared at contact times > 120 s (**Figure S7b,c, Supporting Information**). pKO-β1 fibroblasts established similar adhesion force to FNIII7-10- functionalized 12.8 kPa as to 55.5 kPa PAA gels, while they established lower adhesion force to 0.1 kPa and 4.7 kPa PAA gels. Compared to the adhesion force to 55.5 kPa PAA gels, pKO- β1 fibroblasts established 2- to 6-fold lower adhesion force to 0.1 kPa PAA gels at ≤ 60 s contact times. While we did not observe significant differences between the adhesion force of pKO-β1 fibroblasts to 0.1 kPa and 4.7 kPa PAA gels or to 12.8 kPa and 55.5 kPa PAA gels, we observed a tendency towards lower adhesion force to softer gels. Especially at 360 s contact time, pKO-β1 fibroblasts adhering to 0.1 kPa PAA gels populated only low adhesion forces, while 30% of the pKO-β1 fibroblasts adhering to 4.7 kPa PAA gels populated medium adhesion forces (**Figure S7d,e, Supporting Information**). Hence, we analyzed the adhesion strengthening of pKO-β1 fibroblasts to the FNIII7-10-functionalized PAA gels (**Figure 4b**). pKO-β1 fibroblasts strengthened the adhesion at a similar rate to 55.5 kPa and 12.8 kPa PAA gels, which was two- and four-fold higher compared to 4.7 kPa and 0.1 kPa PAA gels, respectively. Interestingly, the adhesion strengthening rate to 4.7 kPa PAA gels was ∼1.7-fold higher compared to 0.1 kPa PAA gels, which reflected the trend towards lower adhesion forces to the softest PAA gel.

To understand whether the adhesion strengthening mode switch is involved in the mechanosensitive adhesion initiation regulation, we quantified the adhesion force of pKO-β1 fibroblasts to FNIII7-10mSyn functionalized 55.5 kPa and 0.1 kPa PAA gels (**Figure 4c**). The inability to bind to the synergy site reduced the adhesion force of pKO-β1 fibroblasts to FNIII7- 10mSyn functionalized 55.5 kPa PAA gels at 5, 20, 240 and 360 s contact time. However, pKO-β1 fibroblasts established similar adhesion force to FNIII7-10mSyn coated glass and to FNIII7-10mSyn functionalized 55.5 kPa PAA gels (**Figure S7a, Supporting Information**). Moreover, the adhesion of pKO-β1 fibroblasts to FNIII7-10 functionalized 0.1 kPa PAA gels was independent of their ability to bind to the synergy site. Interestingly, we observed a clear trend that pKO-β1 fibroblasts establish lower adhesion forces to 0.1 kPa than to 55.5 kPa FNIII7-10mSyn functionalized PAA gels with increasing contact time. Hence, we also quantified whether the adhesion strengthening of pKO-β1 fibroblasts depended on the stiffness of the FNIII7-10mSyn functionalized PAA gels (**Figure 4d**). Indeed, the adhesion strengthening rate of pKO-β1 fibroblasts to 55.5 kPa PAA gels was 2-fold higher than to 0.1 kPa PAA gels, showing that even if pKO-β1 fibroblasts were unable to bind to the synergy site, they could still respond to the stiffness of FNIII7-10mSyn.

We next investigated whether pKO-αv fibroblasts were able to respond to the FN stiffness and quantified their adhesion force to FNIII7-10 functionalized 55.5 kPa and 0.1 kPa PAA gels (**Figure 4c**). While the adhesion force of pKO-αv fibroblasts to these vastly differently stiff PAA gels was not statistically significant different, we also observed a clear tendency towards lower adhesion force to 0.1 kPa PAA gels. Hence, we quantified whether the adhesion strengthening rate of pKO-αv fibroblasts depended on the functionalized gel stiffness (**Figure 4e**). Similar to pKO-β1 fibroblasts adhering to differently stiff FNIII7-10mSyn functionalized PAA gels, we also observed a 1.6-fold higher adhesion strengthening rate to 55.5 kPa PAA gels compared to 0.1 kPa PAA gels, showing that also for pKO-αv fibroblasts the adhesion initiation to FNIII7-10 was mechanosensitive.

In summary, pKO-β1 and pKO-αv fibroblasts respond to the stiffness of their FN environment in the absence of actomyosin contractility. However, only pKO-β1 fibroblasts triggered a mechanosensitive response to FN substrates stiffer than 5 kPa, which increased their adhesion drastically within seconds.

## Conclusion

Here, we investigated how actomyosin contractility-mediated mechanosensing regulates adhesion initiation of fibroblasts to FN substrates having different stiffnesses by employing either α5β1 or αvβ3 integrins. This adhesion regulation occurs long before integrin clusters become optically visible. On stiff FN substrates of >5 kPa, we observe fibroblasts adhering *via* α5β1 integrin to switch from a slow adhesion strengthening to a fast adhesion strengthening mode within the first 360 s of contact time (**Figure 5**). However, the fibroblasts remain in the slow adhesion strengthening mode on soft FN substrates (<5 kPa). Fibroblasts adhering to FN *via* αvβ3 integrin primarily strengthen adhesion slowly, and independently of myosinII, within 360 s of contact irrespective of the substrate stiffness. The significant differences in adhesion initiation dynamics of α5β1 or αvβ3 integrins highlight that adhesion initiation is an integrin-type specific process^[52–54]^. Since α5β1 integrin binds FN slower than αvβ3 integrin^[13,42]^ and unbind at similar forces from FN in our experimental conditions, we conclude that the fast adhesion strengthening of α5β1 integrin arises from a rapid intracellular avidity regulation within seconds of ligand binding. The dynamics of the intracellular adhesion regulation is specific for individual cells leading to an increased spread of adhesion forces, especially at longer contact times. Consistent with our findings that adhesion initiation of fibroblasts *via* α5β1 integrin is regulated by intracellular signaling within seconds, we have previously observed that this α5β1 integrin-mediated adhesion initiation can be accelerated by an integrin-crosstalk^[42]^, G-protein coupled receptor signaling^[55]^, externally applied mechanical load^[15]^, and syndecan signaling^[47]^.

**Figure 5.**
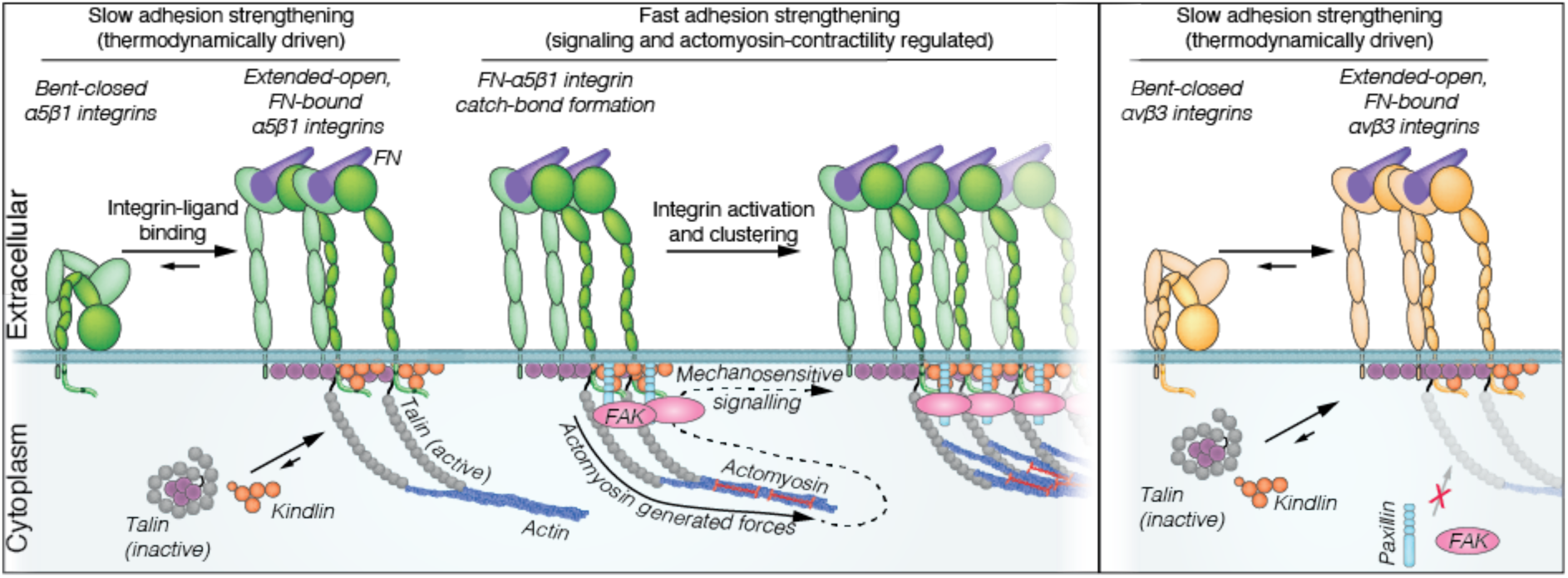
α5β1 integrins but not αvβ3 integrins employ different adhesion strengthening modes to respond to FN stiffness. α5β1 integrins sense the mechanical stiffness of FN during adhesion initiation and eventually switch adhesion strengthening modes of fibroblasts. In fibroblasts, α5β1 integrin employ a slow and a fast adhesion strengthening mode. The slow adhesion strengthening mode presumably depends on the thermodynamic activation of integrins and their maintenance in the ligand bound state by kindlin and talin. This adhesion strengthening mode is only weakly mechanosensitive. Upon myosinII-dependent contractility of the actomyosin cortex, a mechanosensitive signaling pathway is triggered that involves the FN-α5β1 integrin catch bond, paxillin, and FAK. This pathway activates additional integrins and promotes integrin clustering to strengthen cell adhesion to stiff substrates. This mechanosensitive adhesion strengthening mode, dominates the actomyosin- independent response of fibroblasts to FN stiffness. Contrary, αvβ3 integrins cannot switch their adhesion strengthening mode. Within the first 360 s of adhesion initiation, αvβ3 integrin-mediated adhesion strengthening does not depend on integrin-actin engagement. However, also αvβ3 integrin-mediated adhesion initiation is weakly mechanosensitive.

The slow α5β1 integrin adhesion strengthening mode does not depend on the FN synergy site, and therefore not on the catch bond formation of α5β1 integrin (**Figure 5**). This confirms that α5β1 integrin can bind to FN, form adhesion sites, and induce cell spreading irrespective of their binding to the synergy site^[16]^. However, if α5β1 integrin cannot bind to the synergy site fibroblasts show severe adhesion strengthening defects and thus drastic adhesion maturation defects^[16]^. The independence of the slow adhesion strengthening mode on paxillin and FAK signaling indicates this mode to mainly depend on the thermodynamically-driven binding of the integrin ligand^[56–58]^ and the maintenance of the ligand-bound state by talin and kindlin^[15,22,55,59]^. However, we show that the slow adhesion strengthening mode of α5β1 integrin requires the talin rod domain to engage α5β1 integrin to the actin cytoskeleton indicating that the coupling of talin to actin flow^[43]^ applies low forces to integrins^[36,37,60]^. Moreover, our data suggests that the slow adhesion strengthening mode of α5β1 integrin also involves myosinII-mediated contractility, which might be linked to the strengthening of the integrin-FN bond and/or the mechanosensitive recruitment of adhesome proteins to the adhesion site even in the slow adhesion strengthening mode. Nonetheless, even in the absence of the talin rod domain α5β1 integrin can slowly strengthen fibroblast adhesion to FN indicating that maintaining α5β1 integrin in the active conformation can establish weak cell adhesion without coupling to actin, which is in line with previous reports showing that Mn^2+^- activated α5β1 integrins can form small nascent adhesions in the absence of talin^[22]^.

Remarkably, our results show that α5β1 integrin engages to the contractile actomyosin cortex within seconds of after ligand binding, which accelerates adhesion strengthening drastically, resulting in a ∼6-fold adhesion force increase within 360 s (**Figure 5**). Furthermore, our simulations and SCFS experiments suggest that adhesion regulation in response to the intracellular contractility occurs within the first ∼100 s of initiating adhesion, in which α5β1 integrin considerably increases adhesion force. After this adhesion strengthening, α5β1 integrin returns to a moderate adhesion strengthening mode. We show that the [FN-α5β1 integrin] catch bond-paxillin-FAK axis is central to this fast adhesion strengthening and conclude that the recruitment of paxillin and FAK to newly bound α5β1 integrin is promoted by intracellular actomyosin contractility during adhesion initiation^[22,41,61]^. Since inhibition of FAK phosphorylation at Y397 does not completely abolish the fast adhesion strengthening, it indicates that other mechanisms such as adaptor function(s) of FAK and phase separation-induced integrin clustering^[62]^ might be involved. Indeed, *in vitro* experiments show that the recruitment of paxillin and FAK to talin- and kindlin-bound intracellular tails of β1 integrin induces their phase separation and promotes the clustering of β1 integrin-tails^[63]^, which promotes and stabilizes NAs at the leading edge of migrating cells^[22,64]^. Our results thus extend the function of paxillin and FAK in α5β1 integrin-mediated NA assembly to mechanosensitive signaling that regulates α5β1 integrin activity. Thus, the fast adhesion strengthening mode of α5β1 integrin also involves intracellular force-mediated integrin activation, recruitment, and clustering, similar to a previously described instant adhesion regulation of α5β1 integrins to extracellular forces^[15]^. These results highlight that α5β1 integrin instantly senses intracellular and extracellular forces, which triggers active cellular responses. Interestingly, upon initiating adhesion to FN, αvβ3 integrin only marginally involves myosinII contractility within the first 360 s (**Figure 5**). Our results further suggest that within the first 6 min of adhesion initiation the majority of αvβ3 integrins do not require the engagement to the actomyosin cortex by the talin rod domain, which is in line with earlier reports show that αvβ3 integrins are activated and clustered in the presence of the talin head domain^[65]^. Since αvβ3 integrin is not engaged to the contractile actomyosin cortex within the first 360 s it is not surprising that the adhesion initiation of αvβ3 integrin is independent of paxillin recruitment in this timeframe. However, our results indicate that the sparsely populated medium adhesion force population requires αvβ3 integrin to couple to the contractile actomyosin cortex and recruit paxillin. Since αvβ3 integrin requires myosinII-contractility for focal adhesion maturation^[41]^, these results suggest αvβ3 integrin is subjected to actomyosin-mediated contractility and intracellular adhesion strengthening regulation but at much later stages of adhesion initiation than α5β1 integrin. Furthermore, we show that αvβ3 and α5β1 integrins that are not engaged to the actomyosin cortex initiate adhesion similarly, showing that the connection to the actomyosin cortex is essential for the integrin-type specific dynamics in adhesion initiation. These results highlight that the dynamics and likely also the signaling pathways regulating adhesion initiation are integrin-type specific properties.

We here established the combination of SCFS and functionalized PAA hydrogels ranging from 0.1 to 55 kPa, which mimics the stiffness range found for native ECMs^[48,50]^, and provide a unique platform to investigate the mechanisms of mechanosensation of fibroblasts during adhesion initiation. Using this setup, we find that fibroblasts employ two distinct mechanotransduction pathways to regulate adhesion to FN substrates having different stiffnesses. We report a correlation between FN stiffness and adhesion initiation of α5β1 integrin to FN, that is independent of the binding to the synergy site and of αvβ3 integrin to FN. Since under these conditions α5β1 and αvβ3 integrins only minimally involve actomyosin contractility, we conclude that this mechanotransduction pathway is dominated mainly by changes of bond lifetimes due to the different stiffness of the substrate^[36,37,66,67]^. However, both integrins marginally strengthening adhesion 1.5- to 2-fold faster to the stiffest compared to the softest FN substrate, despite the nearly 400-fold higher stiffness. This observed correlation between adhesion strengthening and the FN stiffness follows the predictions of the clutch model that integrin mediated adhesion strengthens with increasing stiffness^[36,37]^. However, it should be noted that the molecular clutch model is defined for force loading within focal adhesions in cells that have already spread on an ECM substrate, while models that address the early phases of ECM recognition and the subsequent initiation of cell adhesion remain absent.

The second mechanotransduction pathway regulating adhesion is exclusively triggered by α5β1 integrin within the first 360 s of contact to FN and requires actomyosin contractility and signaling. We show that α5β1 integrin employ their adhesion strengthening mode switch described above to respond to the stiffness of their environment within seconds of establishing adhesion. While fibroblasts expressing α5β1 integrins remain in the low adhesion strengthening mode on soft FN (<5 kPa), they can switch into the high adhesion strengthening mode, and hence show drastically higher adhesion on stiff FN (>5 kPa). The actomyosin-mediated adhesion switch results in an up to ∼6-fold higher adhesion forces from the stiffest (∼55 kPa) to the softest FN (∼0.1 kPa). However, α5β1 integrin must form a catch bond with FN to switch into the fast adhesion strengthening mode. Since adhesion initiation of α5β1 integrin to stiff FN (>5 kPa) depends on α5β1 integrin binding to the synergy site in FN, but the adhesion to soft FN (<5 kPa) is independent of the engagement of the synergy site, shows that the catch bond of α5β1 integrin is a central feature to sense and respond to stiff FN. Our results further highlight that the active cellular response (signaling) to the stiffness during adhesion initiation dominates the myosinII independent response. Hence, our results extend the current clutch model describing the talin unfolding-driven adhesion maturation during the first hours of cell attachment, on FN substrates stiffer than 5 kPa^[35,36]^, by showing that within comparable stiffness regimes, α5β1 integrin employs a mechanosensitive signaling hub that activates and clusters unbound α5β1 integrins to accelerate adhesion initiation long before adhesion sites are optically visible. Moreover, since only α5β1 integrin but not αvβ3 integrin responds to the FN stiffness of by myosinII-mediated contractility and signaling, we show that adhesion strengthening and mechanotransduction during adhesion initiation are integrin specific properties.

## Methods

### Cell Culture

pKO-β1^[41]^, pKO-αV^[41]^, pKO-β1 pxn KO^[47]^, and pKO-αV pxn KO fibroblasts were maintained in DMEM Glutamax (Gibco-Life technologies) supplemented with 100 units ml^−1^ penicillin (Gibco-Life technologies), 100 μg ml^−1^ streptomycin (Gibco-Life technologies) and 10% (v/v) fetal calf serum (FCS, Sigma) on FN-coated tissue culture flasks.

### Cell line engineering

To deplete paxillin in pKO-αV fibroblasts, CRISPR/Cas9-based strategy used. Single-guide RNAs (sgRNAs) for two different exons were designed using CHOPCHOP (https://chopchop.cbu.uib.no) for paxillin in Mus musculus genome (mm10/GRCm39). sgRNAs were selected based on higher rank, low self-complementarity, low off-target score, and efficiency score. The sgRNAs (with underlined PAM sequences; 5′-NGG-3′) that we designed targeted two separate exons on paxillin. For paxillin, exon 2 (chr5:115 544 490) [5′- CGTGCCATTGAGGGCCTCGCTGG-3′] and exon 8 (chr5:115 552 099) [5′- GTAAGGTCGTGACCGCCATGGGG-3′] were targeted. After 48 h of transfection, GFP and BFP double positive cells were sorted using a FACS sorter (SONY MA900) and cultured on FN-coated culture dishes. Loss of paxillin was confirmed by western blotting using a monoclonal anti-paxillin antibody (1:2500, abcam, ab32084) with glyceraldehyde 3-phosphate dehydrogenase (GAPDH, 1:2500, Cell Signaling Technology, clone 14C10; 2118S) as loading control. Both primary antibodies were detected by using a horseradish peroxidase (HRP)- conjugated goat anti rabbit secondary antibody (1:2500, BioRad, 170–6515).

### Cantilever and substrate preparation

Cantilevers (NPO-D, Bruker) were plasma cleaned (DC-32G; Harrick Plasma) for ∼5 min prior to overnight coating with 2 mg ml^−1^ concanavalinA (ConA, Merck) in PBS at 4° C. For glass surface coating four-segment PDMS masks^[53]^ were attached to glass bottom petri dishes (WPI). Prior to coating glass surfaces were cleaned by thoroughly washing with ethanol and water. Subsequently, the segmented glass surfaces were coated overnight with 20 µl FNIII7- 10, FNIII7-10ΔRGD or FNIII7-10mSyn (all 50 µg ml^−1^ in PBS) at 4° C. For ligand density characterization of easycoat polyacrylamide (PAA) gels (Softview) were functionalized according to manufacturer guidelines with 50 µg ml^−1^ bovine FN (in PBS) for 30 min at room temperature (RT). For FN-ligand density characterization and for SCFS on FNIII7-10-density adjusted PAA gels, PAA gels were functionalized with 500 µl PBS supplemented with different ratios of 50 µg ml^−1^ FN and 50 µg ml^−1^ BSA or 50 µg ml^−1^ FNIII7-10 and 50 µg ml^−1^ FNIII7- 10ΔRGD, respectively, for 30 min at RT. FN:BSA or FNIII7-10:FNIII7-10ΔRGD ratios were 1:1, 3:2, 7:3, and 1:0 for 100 kPa, 25 kPa, 8 kPa, and 0.2 kPa (all nominal) PAA gels, respectively.

### Single-cell force spectroscopy (SCFS)

For SCFS, a AFM (Nanowizard II) having a CellHesion module and a PetriDish Heater (all JPK, Berlin, Germany) to provide 37°C ambient temperature was mounted on an inverted optical microscope (ObserverZ1, Zeiss, Germany). V-shaped, 200 µm long NPO-D cantilevers having nominal spring constants of 0.06 N m^−1^ were used for SCFS. The sensitivity of the AFM setup was determined on glass surfaces and the cantilevers’ spring constant was calibrated by the thermal noise method prior to their use in SCFS.

Fibroblasts were grown to a confluency of ∼ 80%, serum-starved overnight, trypsinized (0.25% trypsin/EDTA; Gibco) for 2 min at 37°C and resuspended in SCFS media (DMEM supplemented with 20 mM HEPES) containing 1% FCS. Fibroblasts were allowed to recover from trypsinization for at least 30 min prior to SCFS experiments^[68]^. The fibroblasts suspension was pipetted onto the Petri dish, the ConA-coated cantilever was positioned above a single rounded fibroblast to which it was approached with a speed of 10 µm s^−1^ until a force of 5 nN was detected. The cantilever height was maintained constant for 5 s before retracting the cantilever for >50 µm. Thereafter, the cantilever-bound fibroblast was allowed to firmly bind to the cantilever for 3 – 5 min before conducting SCFS. Subsequently, the fibroblast attached to the cantilever was approached to the substrate at 5 µm s^−1^ until reaching the contact force of 1 nN. The cantilever height was maintained constant for 5, 20, 60, 120, 240 or 360 s, before retracting the cantilever at 5 µm s^−1^ for >90 µm to fully detach the fibroblast from the substrate. This experimental cycle was repeated for a single cantilever-bound fibroblast for all contact times or until morphological changes (i.e., cell spreading) was observed. Before starting a new adhesion force measurement cycle, the cantilever-bound fibroblast was allowed to recover for at least the contact time of the measurement and the location on the substrate was changed. The order of contact times characterized was randomized. Most fibroblasts spread on the cantilever before we could measure their adhesion forces for all contact times and thus had to be replaced by fresh round fibroblasts to continue the adhesion measurements.

For SCFS with inhibitors, myosinII-mediated contractility was inhibited by 20 µM blebbistatin (Merck) and FAK phosphorylation was inhibited by 10 µM Y11 (Torcis). Fibroblasts detached from the culture dish were incubated with the given concentration of inhibitors for 30 min prior and throughout SCFS.

#### Bootstrapping

To simulate the adhesion strengthening of individual fibroblasts, we utilized Python 3.9.6 to bootstrap the dataset containing cell adhesion forces at contact times of 5, 20, 60, 120, 240 or 360 s, resulting in 4000 time series representing potential single cell adhesion strengthening scenarios. This process involved employing the sampling function with sample replacement from the Pandas v.2.0 library. For each adhesion force time series, we randomly sampled data points from the adhesion force pool while satisfying two conditions: Compared to the previous contact time, a cell either (1) increases adhesion, or (2) maximally decreases adhesion by 10%. These conditions were implemented by masking the Pandas DataFrame containing the whole adhesion data with an appropriate filter for each time step and time series.

### Stiffness measurements

For stiffness measurements of FN fragment functionalized PAA gels, an NPO-D cantilever with a 6 µm diameter silica bead (Kisker Biotech GmbH) attached to its apex was used. Prior to stiffness measurements, the sensitivity of the AFM setup was determined on a glass surface prior to determining the spring constant of the cantilever by the thermal noise method. For indentation experiments, the beaded cantilever was positioned above the FNIII7-10-coated PAA gel. A grid of 2 x 2 positions within 100 µm x 100 µm was defined for indentation experiments. The beaded cantilever was approached to the PAA gel at 5 µm s^−1^ until recording a force of 1 nN. Subsequently, the cantilever was retracted and moved to the next position in the grid. During each approach and retraction cycle of the cantilever a FD curve was recorded. Five different regions per PAA gel and four independently prepared gels were tested per stiffness value of the gel.

### Immunofluorescence

FN-functionalized PAA gels were washed twice with PBS and blocked with PBS containing 3% BSA for 30 min at RT. PAA gels were incubated with a 1:100 dilution of FN antibodies (ab2413, abcam) in PBS containing 3% (w/v) BSA for 1 h at RT. After 3 times washing with PBS, PAA gels were incubated with secondary anti-rabbit IgG antibody conjugated with Alexa488 (ab150077, abcam, 1:200 in PBS containing 3% (w/v) BSA). Images were acquired with an inverted microscope (ObserverZ1, Zeiss) equipped with an LSM700 confocal point scanner (Zeiss) and a 25x/0.8 water immersion objective (LD LCI Plan-Apochromat, Zeiss). To control for specific fluorescent signal, a smaller region was photobleached by imaging for up to 20 times with high laser power. Subsequently, a larger area was imaged with low laser power.

For fibroblasts spreading experiments, PAA gels were functionalized with FNIII7-10 to have similar RGD densities (see *Cantilever and substrate preparation*). Subsequently, pKO- β1 fibroblasts were seeded for 90 min on the functionalized PAA gels before fixing them with 4% PFA (Sigma) in PBS for 15 min at RT. After thorough washing with PBS, fibroblasts were permeabilized with 0.2% (v/v) Triton X-100 in PBS for 5 min at RT. Samples were washed with excess PBS and blocked with PBS containing 3% (w/v) BSA and 0.1% (v/v) Tween-20 (PBT) for 30 min at RT. Subsequently, fibroblasts were incubated with a 1:400 dilution of an anti-paxillin AB (ab32084, abcam) in PBT for 1 h at RT. After 3 times washing with PBS for 5 min at RT, samples were incubated with anti-rabbit IgG antibody conjugated with Alexa647 (ab150079, abcam, 1:200), phalloidin-Alexa488 (Invitrogen, 1:400) and Hoechst (1:1000) all in PBT for 1 h at RT. Images were acquired using an inverted microscope (ObserverZ1, Zeiss) equipped with an LSM700 confocal point scanner (Zeiss) and a 40x/0.75 objective (EC Plan-Neofluar, Zeiss). An image of a single plane containing the paxillin signal was used for spreading area quantification.

### Data Analysis

To determine cell adhesion forces, JPK inbuild routines were used. Retraction FD curves were corrected for baseline and drift, and the maximum downward deflection of the cantilever quantified the adhesion force.

Rupture events were analyzed using an in-house routine (IgorPro8.04) to quantify the forces rupturing single integrin-FN bonds. Tether events were excluded from analysis, as they represent the force required to extract a tether from the cell membrane. Rupture and tether events were distinguished by the slope prior to the unbinding event, where tether plateaus had a maximum angle of ∼ 10° (**Figure S1**). The minimum rupture force analyzed was set to 20 pN and the maximum rupture force to 150 pN. For double Gaussian fits, rupture forces up to 90 pN were considered.

Young’s moduli of PAA gels were determined using the inbuild AFM data analysis software (JPK, Version spm-4.3.55). Recorded approach FD curves were offset- tilt-, and cantilever deflection corrected. Afterwards, the approach FD curve was fitted by the Hertz model with a Poisson ration of 0.5. The contact point in FD curves was determined automatically.

To quantify FN densities of functionalized PAA gels, six line profiles per image were analyzed for fluorescent intensities and averaged. Intensities were normalized to the average intensities of the stiffest PAA gel. For spreading area quantification, an in-house created macro was developed for ImageJ (Version 2.3.0/1.53q). For the segmentation of cell spreading areas from paxillin signals, RenyiEntropy^[69]^ and MinError^[70]^ methods were respectively used.

Statistical analysis was performed using Prism 9 (GraphPad).

## Acknowledgements

We thank R. Fässler for providing essential cell lines, fruitful discussions and critical reading of the manuscript, and G. Ammirati for providing beaded cantilevers. We thank the single cell facility in the Department of Biosystems Science and Engineering, ETH Zürich for their continuous support. This work was supported by the Swiss National Science Foundation (SNF; grants no. 31003A_182587/1 and 310030_215690/1).

## Author Contributions

NS and DJM conceived and designed the study. NS performed most of the experiments. US engineered pxn KO fibroblasts. NS and MMN performed SCFS experiments for large adhesion force data sets. NS analyzed SCFS data. GF performed bootstrapping. NS analyzed bootstrapped data set. JCA assisted in confocal imaging and developed the algorithm for cell spreading area analysis. NS and DJM evaluated experimental progress and data. All authors discussed the experiments and wrote the manuscript.

## Table of contents

We find that fibroblasts initiate adhesion using two distinct modes in response to fibronectin stiffness. Both α5β1 and αvβ3 integrins exhibit slow adhesion strengthening on soft substrates, independent of actomyosin contractility. On stiff fibronectin, however, only α5β1 integrins switch to a fast adhesion mode, driven by myosin II and mechanosensitive signaling. This reveals integrin-specific differences in stiffness sensing and adhesion dynamics.

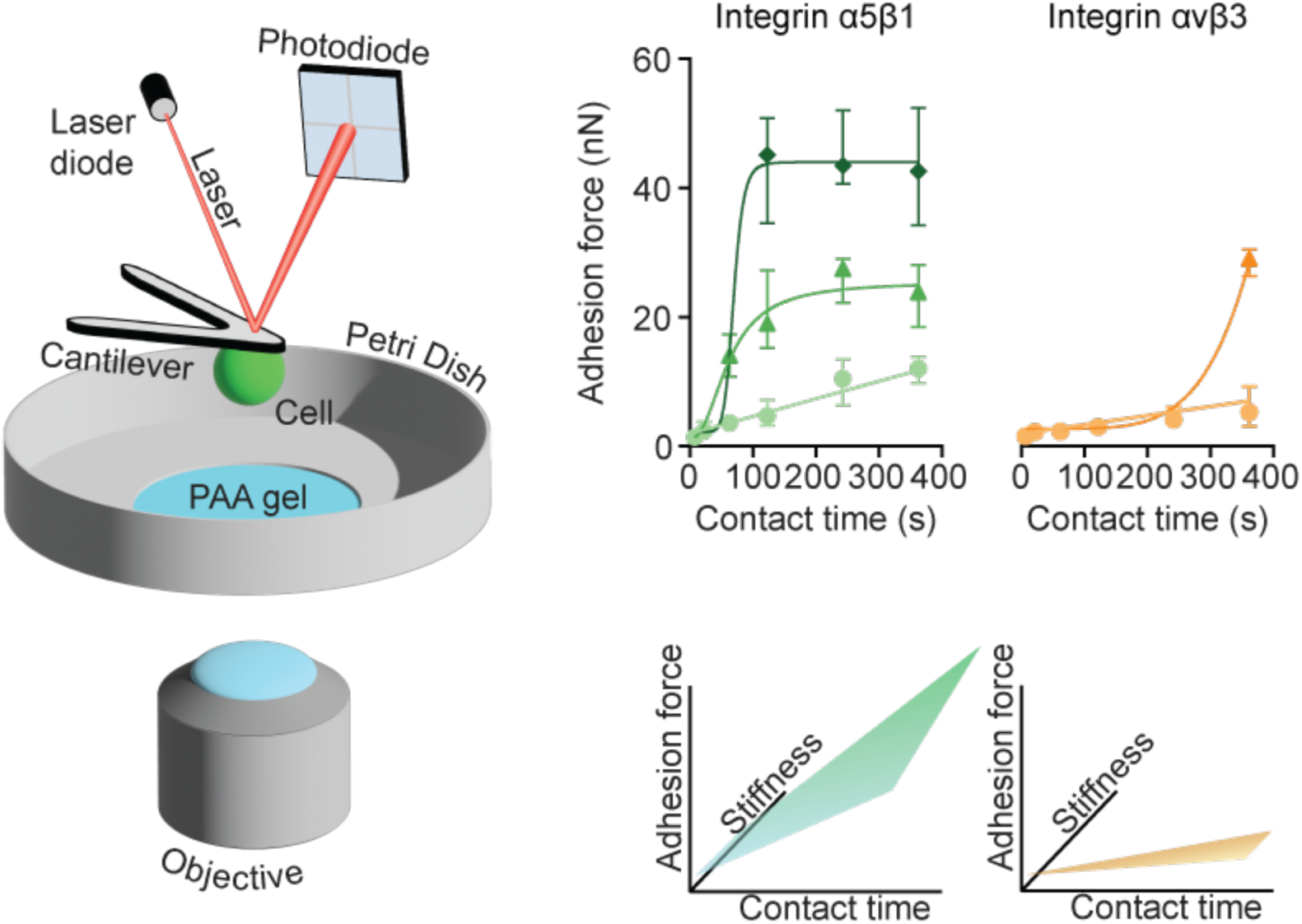

## Supporting Information

**Figure S1.**
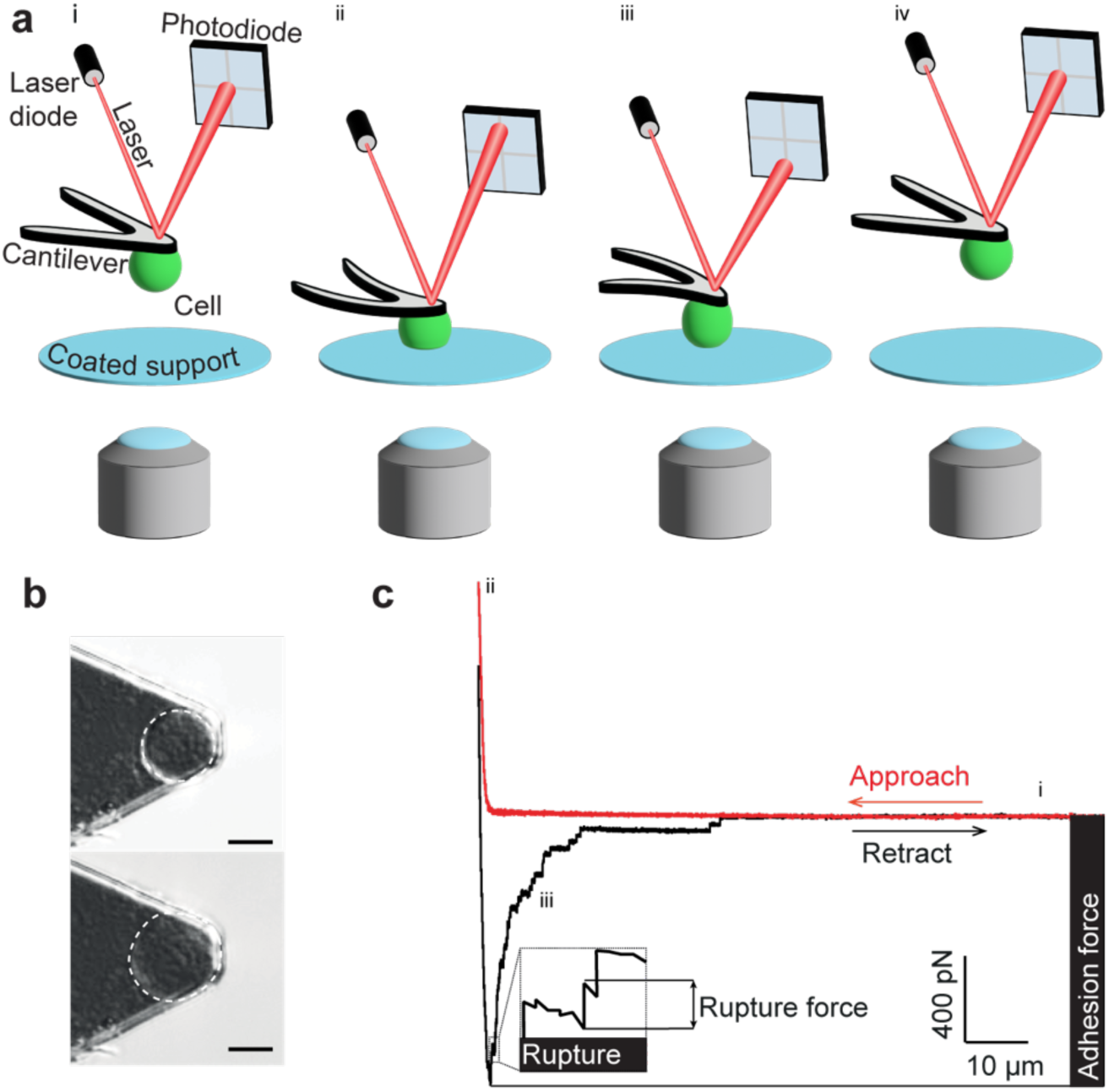
Atomic force microscopy (AFM)-based single-cell force spectroscopy (SCFS) setup to quantify the adhesion force of single fibroblasts. **a)** Schematic illustration of SCFS setup and approach. i) A rounded cantilever-bound fibroblast is brought into contact with the substrate at a preset contact force. ii) Once the contact force is reached, the height of the cantilever is maintained constant, and the cell is allowed to interact with the substrate for a given contact time. iii) The cantilever is retracted iv) until the fibroblast is fully separated from the substrate to measure the adhesion force of the fibroblasts to the substrate. **b)** Differential interference contrast (DIC) image of the AFM cantilever with a single fibroblast attached (top). Upon the observed morphological changes (bottom), the fibroblast is discarded. Scale bar, 10 µm. **c)** Representative force-distance curve of a SCFS experiment, which was recorded upon approaching (red curve) and retracting (black curve) a single fibroblast to and from FNIII7-10 coated glass substrates, respectively. The maximum downward deflection of the cantilever measures the adhesion force, which is the maximum force with which the fibroblast adheres to its substrate. After the adhesion force peak, smaller unbinding events (ruptures) are observable that quantify the maximum force that a single integrin can bear before it unbinds from the ligand.

**Figure S2.**
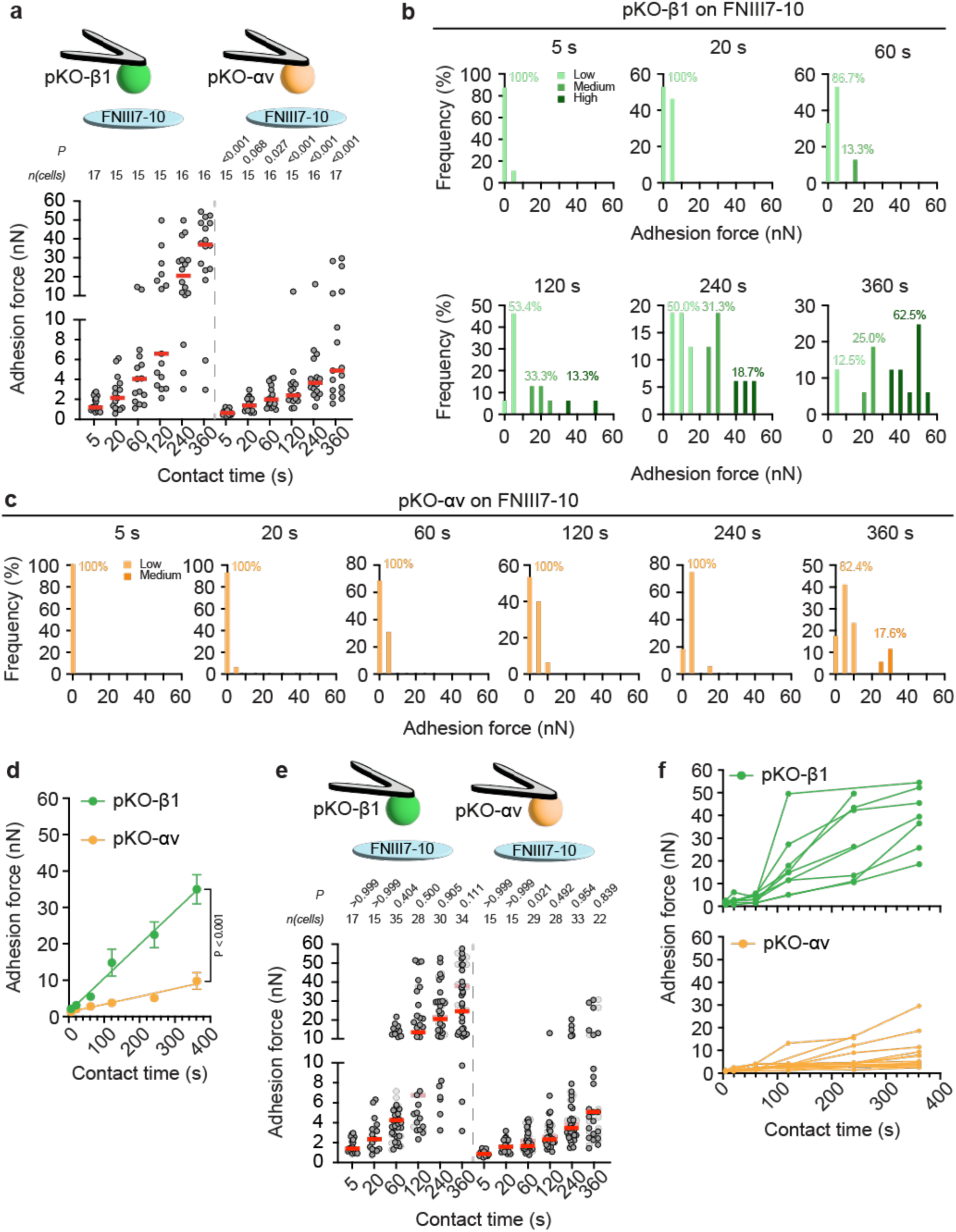
α5β1 but not αvβ3 integrins switch the adhesion strengthening mode to FNIII7-10 within the first minutes of initiating adhesion. **a)** Adhesion forces of (left) pKO- β1 and (right) pKO-αv fibroblasts to FNIII7-10 coated glass surfaces at given contact times. Dots represent adhesion forces of individual fibroblasts, red bars the median, and *n*(cells) the number of fibroblasts tested. *P* values comparing adhesion forces of pKO-β1 and pKO-αv fibroblasts at the respective contact time were calculated using two-sided Mann-Whitney tests. **b,c)** Histograms with a bin width of 5 nN show the adhesion force distribution of b) pKO-β1 and c) pKO-αv fibroblasts adhering to FNIII7-10 for the given contact time. Light color bars indicate the low adhesion force population (<15 nN), medium color bars the medium adhesion force population (15 – 35 nN), and dark color bars the high adhesion force population (>35 nN). Numbers indicate the occupancy (in %) of each population. **d)** Adhesion strengthening of the whole pKO-β1 or pKO-αv fibroblast population was quantified as a slope of a linear function (lines) fitted to all adhesion forces at all contact times. Dots represent the mean adhesion force and bars the standard error of the mean (SEM). The given *P* value compares the linear fit of both data sets and was calculated by extra sum-of-squares F-tests. **e)** Large adhesion force data sets were acquired for contact times ≥ 60 s to quantify adhesion strengthening of subpopulations. The original data set is given in the background for comparison of adhesion force distribution. Dots represent adhesion forces of individual fibroblasts, red bars the median, and *n*(cells) the number of fibroblasts tested. *P* values comparing given and reference data were calculated using two-sided Mann-Whitney tests. **f)** Adhesion strengthening of individual pKO-β1 (top) or pKO-αv (bottom) fibroblast as detected by SCFS (data taken from **a**).

**Figure S3.**
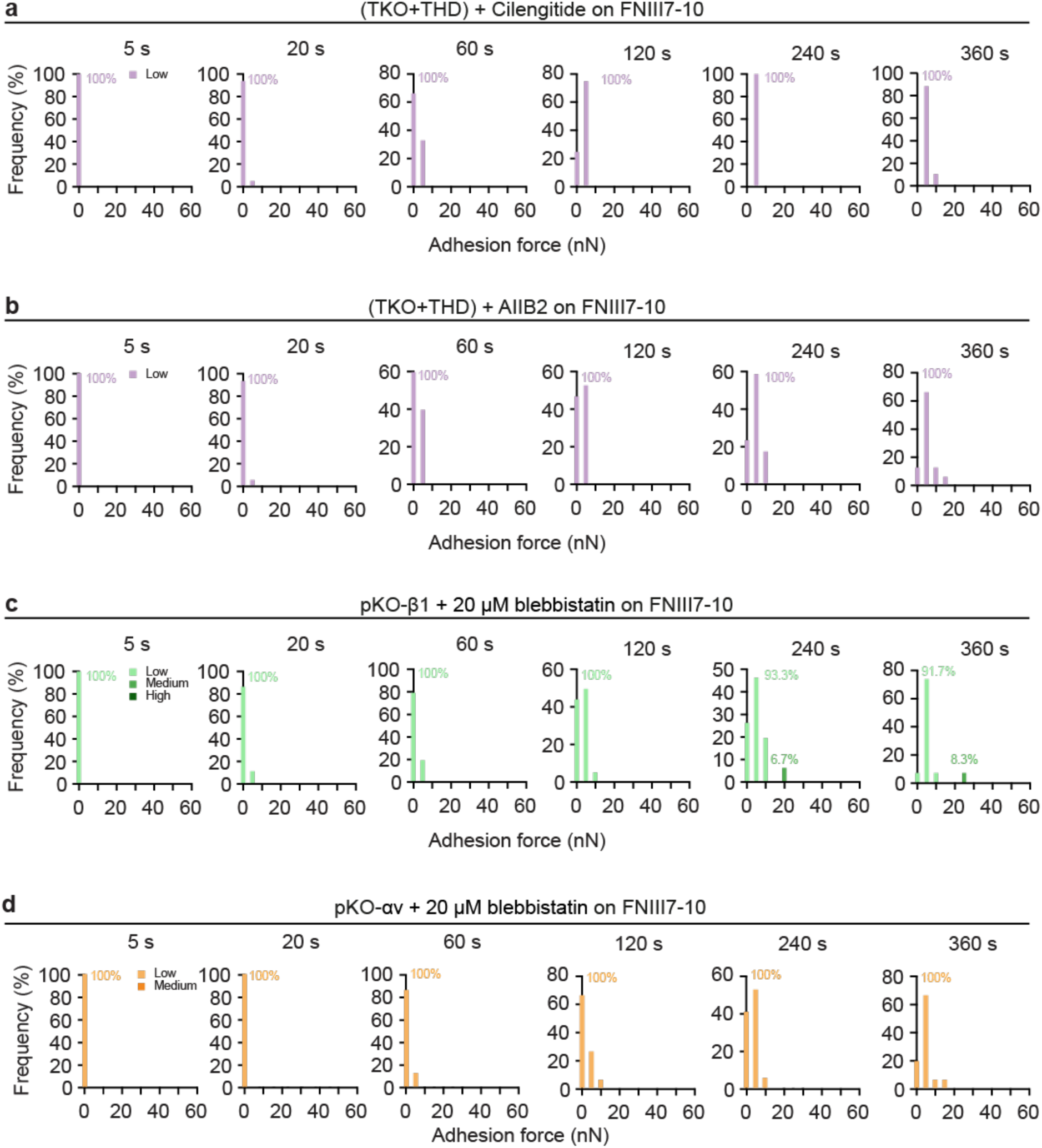
Medium and high adhesion force populations depend on the engagement of integrins to the contractile actomyosin. Adhesion force distribution of (a,b) talin1/2- depleted and talin1 head domain re-expressing (TKO+THD) fibroblasts in the presence of (a) cilengitide or (b) AIIB2 and (c,d) myosinII-contractility inhibited (c) pKO-β1 or (d) pKO-αv fibroblast. Histograms with a bin width of 5 nN show the adhesion force for the given contact time. The color of the bars indicate the adhesion force population as annotated. The numbers indicate the occupancy of the populations.

**Figure S4.**
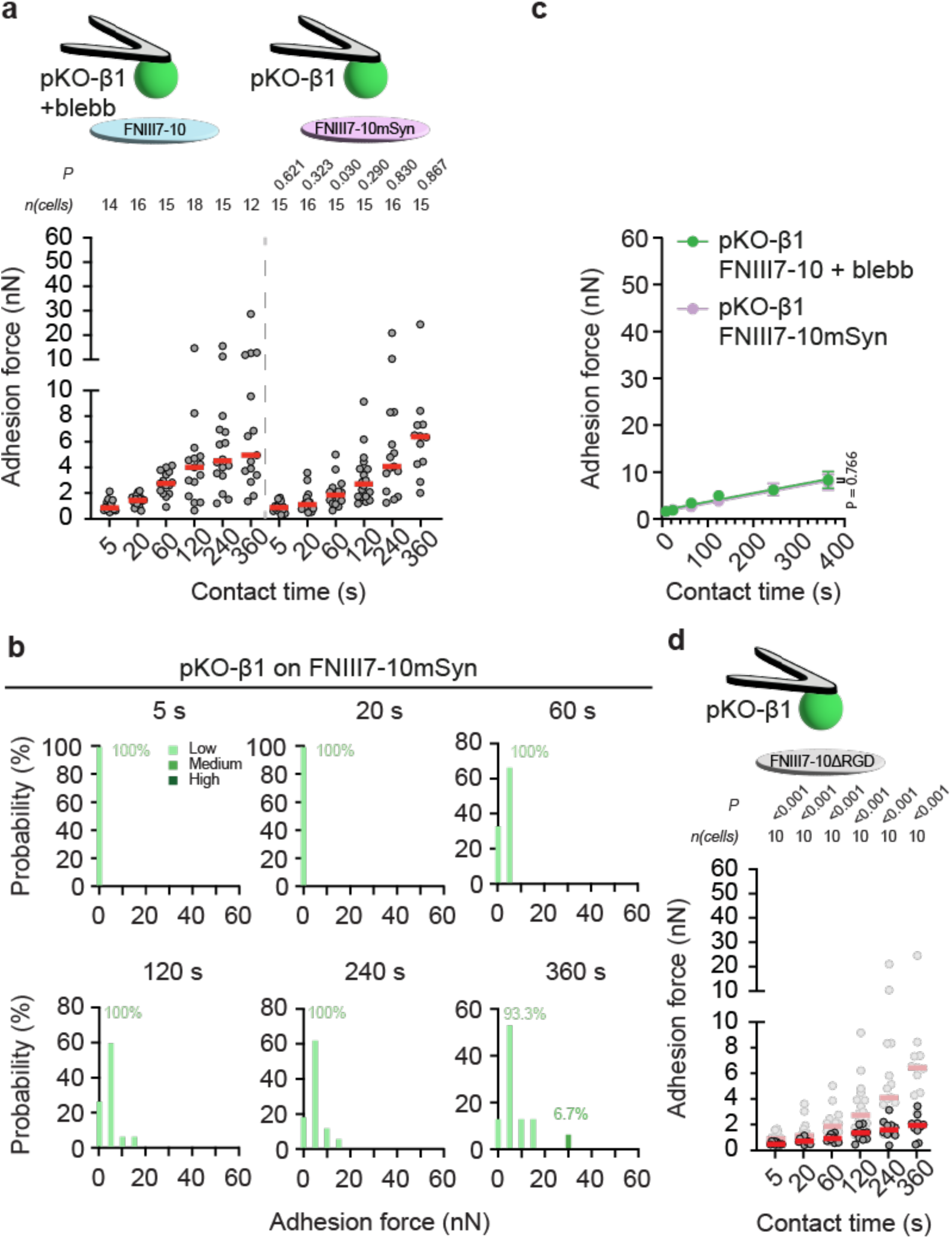
The fast adhesion strengthening mode of pKO-β1 fibroblasts depends on the FN synergy site. **a)** Adhesion forces of (left) 20 µM blebbistatin-treated pKO-β1 fibroblasts to FNIII7-10 (data taken from Fig. 2d) and (right) of untreated pKO-β1 fibroblasts to FNIII7-10mSyn (data taken from Fig. 3a). Dots represent adhesion forces of individual fibroblasts, red bars the median, and *n*(cells) the number of fibroblasts tested. *P* values comparing adhesion forces at the respective contact time were calculated using two-sided Mann-Whitney tests. **b)** Histograms with a bin width of 5 nN show the adhesion force distribution of pKO-β1 fibroblasts adhering to FNIII7-10mSyn for the given contact time. The color of the bars indicate the adhesion force population as annotated. The numbers indicate the occupancy of the populations. **c)** Adhesion strengthening of 20 µM blebbistatin-treated pKO-β1 fibroblasts to FNIII7-10 or pKO- β1 fibroblasts to FNIII7-10mSyn was quantified as a slope of a linear function (lines) fitted to all adhesion forces at all contact times. Dots represent the mean adhesion force and bars the standard error of the mean (SEM). The given *P* value compares the linear fit of both data sets and was calculated by extra sum-of-squares F-tests. **d)** Adhesion forces of pKO-β1 fibroblasts to FNIII7-10ΔRGD at given contact times. Adhesion forces of pKO-β1 fibroblasts to FNIII7-10mSyn (data taken from Fig. 3a) are given in grey as reference. Data representation as in a). P values comparing given and reference data were calculated using two-sided Mann-Whitney Tests.

**Figure S5.**
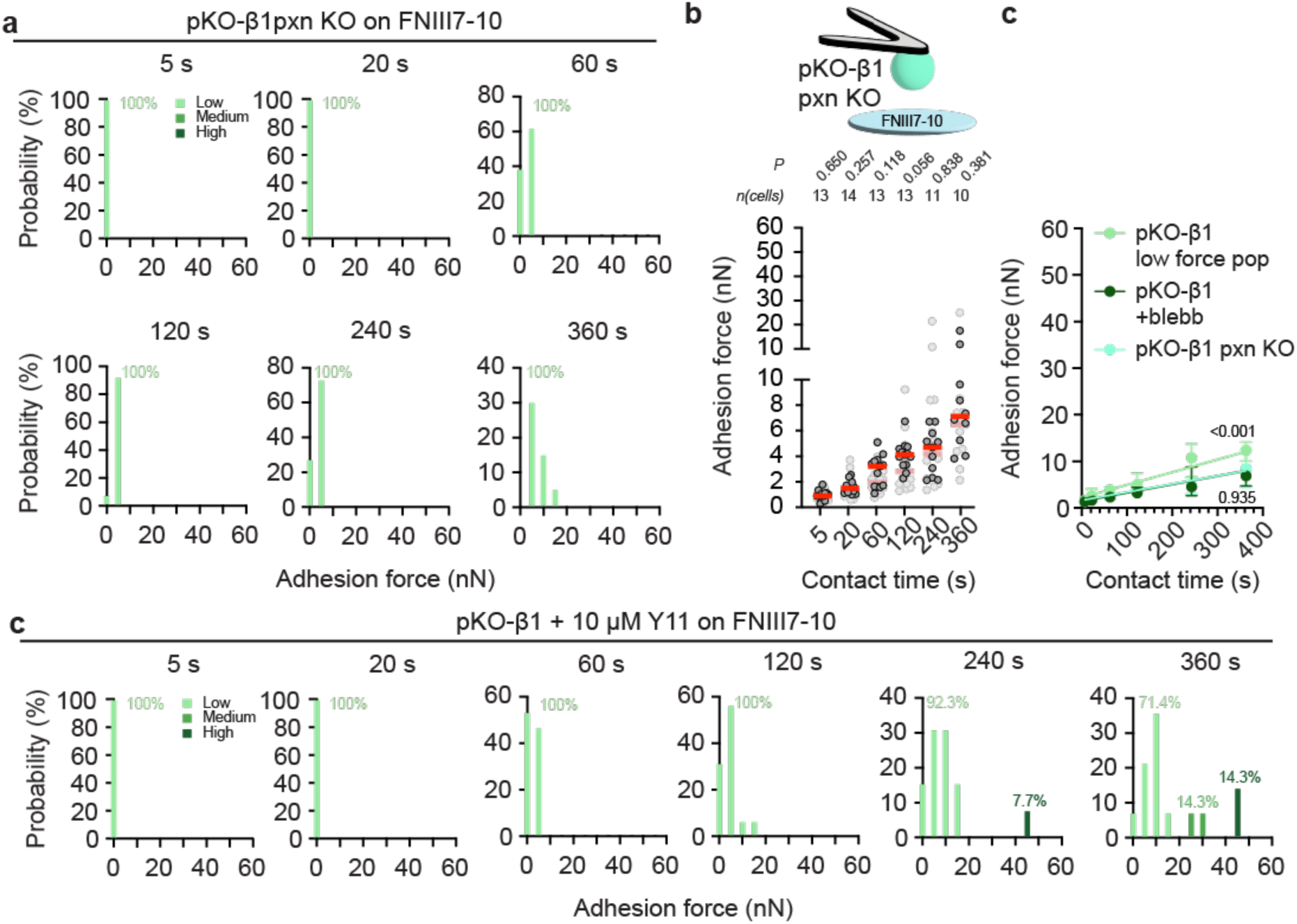
Paxillin and FAK signaling are required for fast adhesion strengthening of pKO-β1 fibroblasts. **a)** Histograms with a bin width of 5 nN show the adhesion force distribution of pKO-β1 pxn KO fibroblasts adhering to FNIII7-10 for the given contact time. Light green bars indicate the low adhesion force population, medium green bars the medium adhesion force population and dark green bars show the high adhesion force population. **b)** Adhesion forces of pKO-β1 pxn KO fibroblasts to FNIII7-10 (data taken from Fig. 5b). Adhesion forces of 20 µM blebbistatin-treated pKO-β1 fibroblasts adhering to FNIII7-10 (data taken from Fig. 4a) are given as grey as reference. Dots represent adhesion forces of individual fibroblasts, red bars the median, and *n*(cells) the number of fibroblasts tested. *P* values comparing adhesion forces at the respective contact time were calculated using two-sided Mann-Whitney tests. **c)** Adhesion strengthening of low adhesion force population pKO-β1 fibroblasts, 20 µM blebbistatin-treated pKO-β1 fibroblasts to FNIII7-10 or pKO-β1 pxn KO fibroblasts to FNIII7-10mSyn was quantified as a slope of a linear function (lines) fitted to all adhesion forces at all contact times. Dots represent the mean adhesion force and bars the standard error of the mean (SEM). The given *P* values compare the adhesion strengthening of (top) the low adhesion force population pKO-β1 fibroblasts to FNIII7-10 with of pKO-β1 pxn KO fibroblasts to FNIII7-10mSyn and (bottom) of the low adhesion force population pKO-β1 fibroblasts to FNIII7-10 and of blebbistatin-treated pKO-β1 fibroblasts to FNIII7-10. The P values were calculated by extra sum-of-squares F-tests. **d)** Histograms with a bin width of 5 nN show the adhesion force distribution of 10 µM Y11-treated pKO-β1 fibroblasts adhering to FNIII7-10 for the given contact time.

**Figure S6.**
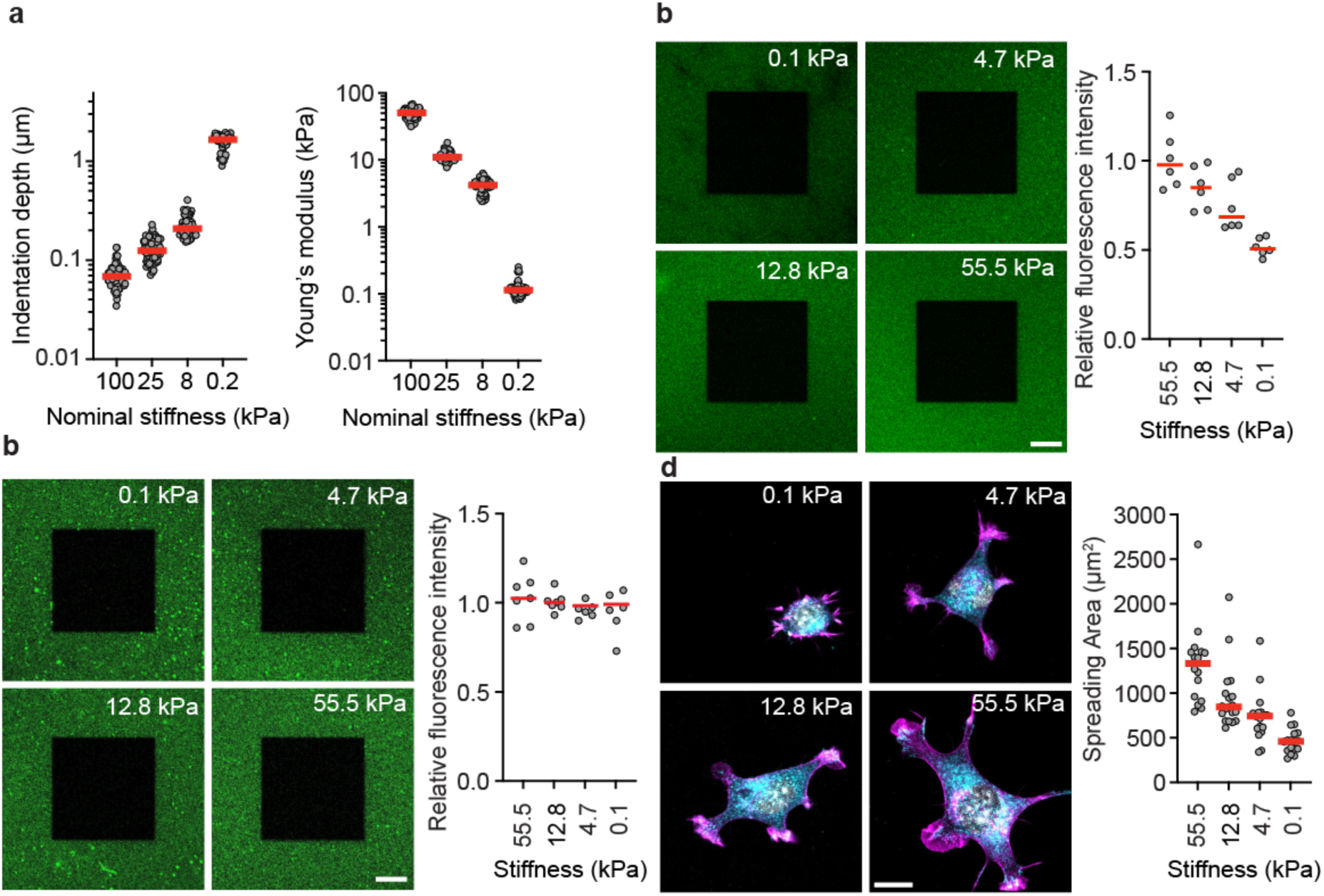
Characterization of PAA gels for SCFS. **a)** Indentation depth (left) and Young’s moduli of PAA gels quantified with a 6 µm glass bead attached to a microcantilever to reach a force of 1 nN. Dots represent indentation depth or Young’s modulus of each experiment and red bars the median. **b)** Left, representative immunofluorescent images of b) FN functionalized PAA gels or c) PAA gels functionalized with adjusted FN concentration having the given stiffness. The dark square in the middle of images shows a photobleached area to validate specific signal. Right, relative fluorescent intensity to the average intensity on the stiffest gel. Dots are the mean relative intensities of six line profiles across each images and red lines their median. **d)** Left, representative images of pKO-β1 fibroblasts spread for 90 min on FNIII7- 10 functionalized PAA gels having the given nominal stiffness. Paxillin staining is shown in cyan, actin staining in magenta and the nucleus in white. Right, quantification of the spreading area of pKO-β1 fibroblasts by paxillin staining on differently stiff substrates. Dots represent spreading areas of individual fibroblasts and red bars the median. All scale bars, 20 µm.

**Figure S7.**
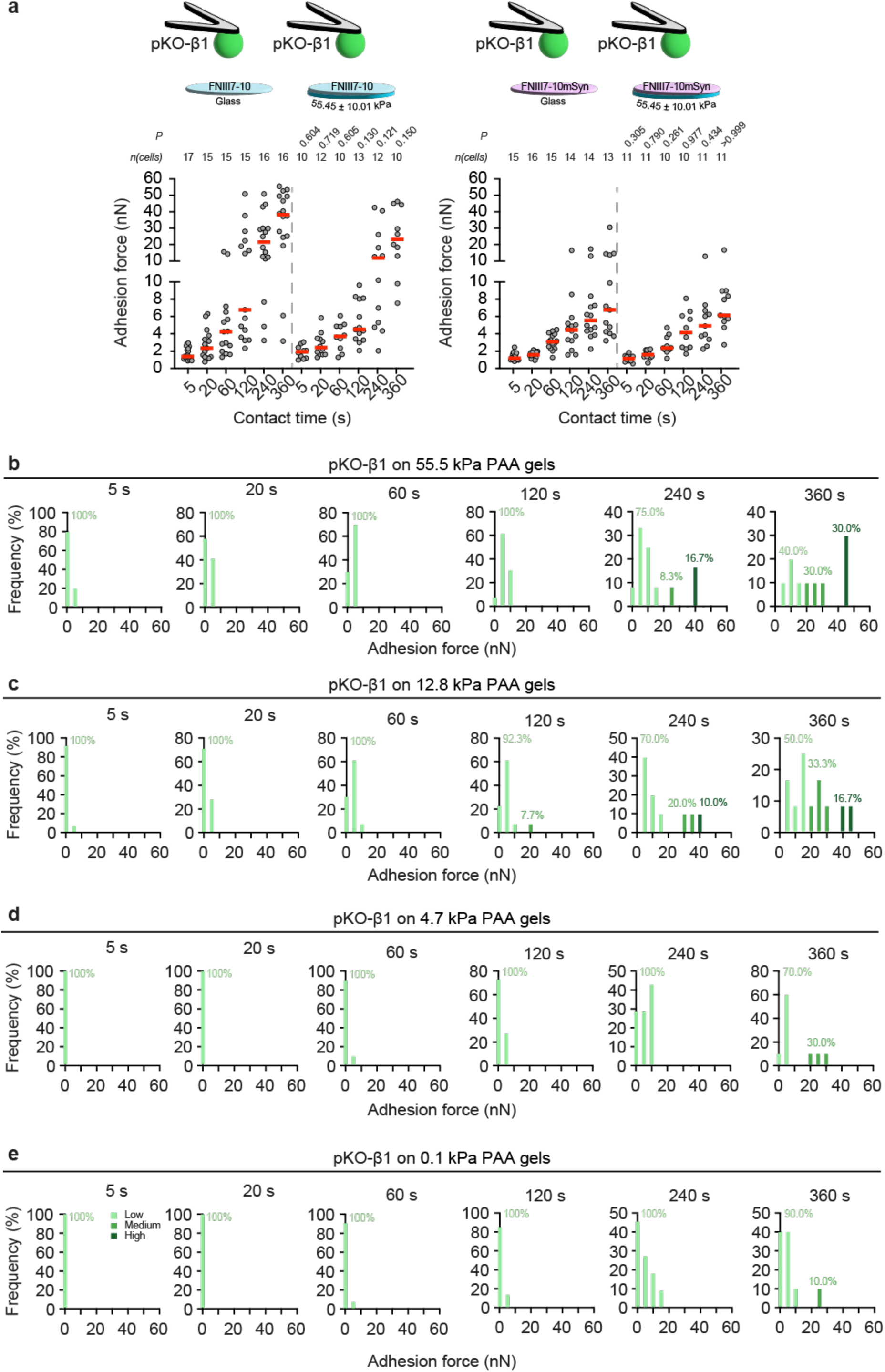
α5β1 integrin populate the low, medium, and high adhesion force population substrate stiffness dependently. a) Adhesion forces of pKO-β1 fibroblasts or adhering to (left) FNIII7-10-coated glass (data taken from Supplementary Fig. 2a) and FNIII7-10- functionalized 55.45 kPa PAA gels (data taken from Fig. 4a) or (right) FNIII7-10mSyn-coated glass (data taken from Fig. 3a) and FNIII7-10mSyn-functionalized 55.45 ± 10.01 kPa PAA gels (data taken from Fig. 4b). Dots represent adhesion forces of individual fibroblasts, red bar the median, and *n*(cells) the number of fibroblasts tested. *P* values comparing adhesion forces at the respective contact time were calculated using two-sided Mann-Whitney tests. **b-e)** Histograms with a bin width of 5 nN show the adhesion force distribution of pKO-β1 fibroblasts adhering to FNIII7-10-functionlaized PAA gels having given stiffness and for the given contact time. Light green bars indicate the low and medium dark green bars the medium adhesion force population. The numbers indicate the occupancy of the populations.

