## Supporting Information for "α5β1 and αvβ3 integrins employ two distinct adhesion strengthening modes to respond to fibronectin stiffness within seconds of initiating adhesion"

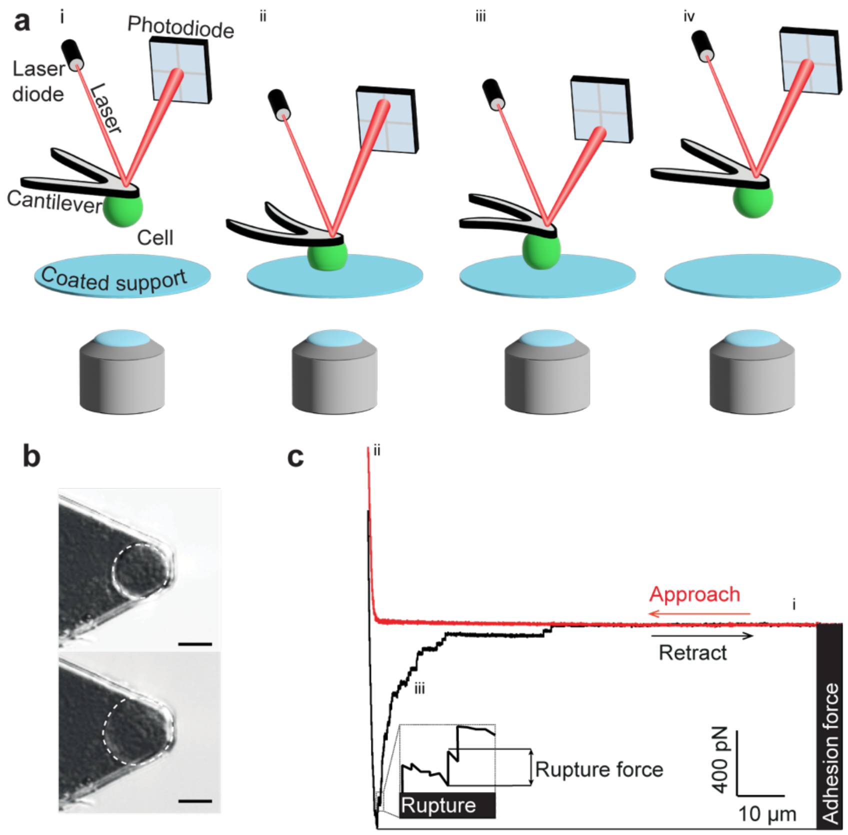


**Figure S1. Atomic force microscopy (AFM)-based single-cell force spectroscopy (SCFS) setup to quantify the adhesion force of single fibroblasts. a)** Schematic illustration of SCFS setup and approach. i) A rounded cantilever-bound fibroblast is brought into contact with the substrate at a preset contact force. ii) Once the contact force is reached, the height of the cantilever is maintained constant, and the cell is allowed to interact with the substrate for a given contact time. iii) The cantilever is retracted iv) until the fibroblast is fully separated from the substrate to measure the adhesion force of the fibroblasts to the substrate. **b)** Differential interference contrast (DIC) image of the AFM cantilever with a single fibroblast attached (top). Upon the observed morphological changes (bottom), the fibroblast is discarded. Scale bar, 10 µm. **c)** Representative force-distance curve of a SCFS experiment, which was recorded upon approaching (red curve) and retracting (black curve) a single fibroblast to and from FNIII7-10 coated glass substrates, respectively. The maximum downward deflection of the cantilever measures the adhesion force, which is the maximum force with which the fibroblast adheres to its substrate. After the adhesion force peak, smaller unbinding events (ruptures) are observable that quantify the maximum force that a single integrin can bear before it unbinds from the ligand.

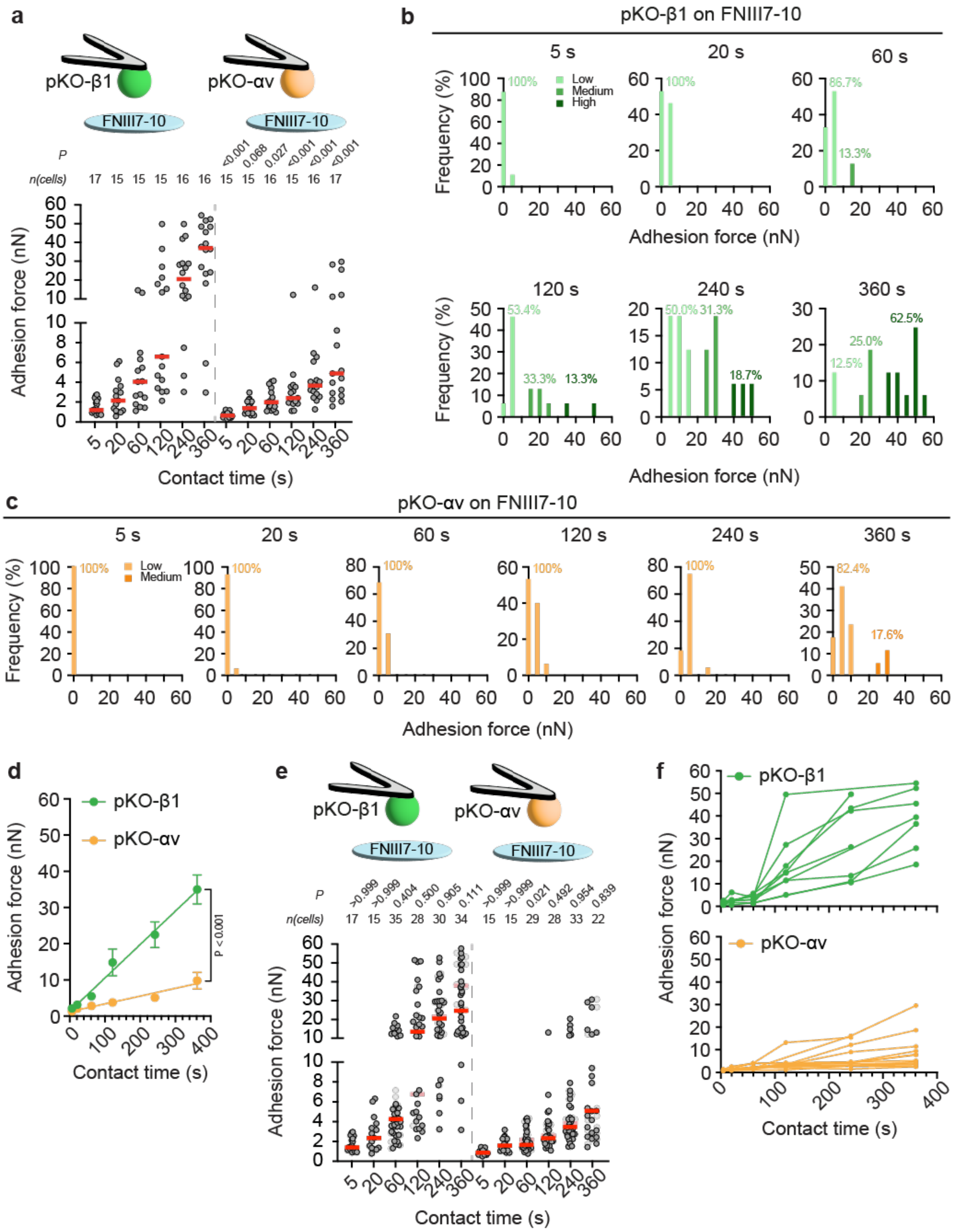


**Figure S2. α5β1 but not αvβ3 integrins switch the adhesion strengthening mode to FNIII7-10 within the first minutes of initiating adhesion. a)** Adhesion forces of (left) pKO-β1 and (right) pKO-αv fibroblasts to FNIII7-10 coated glass surfaces at given contact times. Dots represent adhesion forces of individual fibroblasts, red bars the median, and *n*(cells) the number of fibroblasts tested. *P* values comparing adhesion forces of pKO-β1 and pKO-αv fibroblasts at the respective contact time were calculated using two-sided Mann-Whitney tests. **b,c)** Histograms with a bin width of 5 nN show the adhesion force distribution of b) pKO-β1 and c) pKO-αv fibroblasts adhering to FNIII7-10 for the given contact time. Light color bars indicate the low adhesion force population (<15 nN), medium color bars the medium adhesion force population (15 – 35 nN), and dark color bars the high adhesion force population (>35 nN). Numbers indicate the occupancy (in %) of each population. **d)** Adhesion strengthening of the whole pKO-β1 or pKO-αv fibroblast population was quantified as a slope of a linear function (lines) fitted to all adhesion forces at all contact times. Dots represent the mean adhesion force and bars the standard error of the mean (SEM). The given *P* value compares the linear fit of both data sets and was calculated by extra sum-of-squares F-tests. **e)** Large adhesion force data sets were acquired for contact times ≥ 60 s to quantify adhesion strengthening of subpopulations. The original data set is given in the background for comparison of adhesion force distribution. Dots represent adhesion forces of individual fibroblasts, red bars the median, and *n*(cells) the number of fibroblasts tested. *P* values comparing given and reference data were calculated using two-sided Mann-Whitney tests. **f)** Adhesion strengthening of individual pKO-β1 (top) or pKO-αv (bottom) fibroblast as detected by SCFS (data taken from **a**).

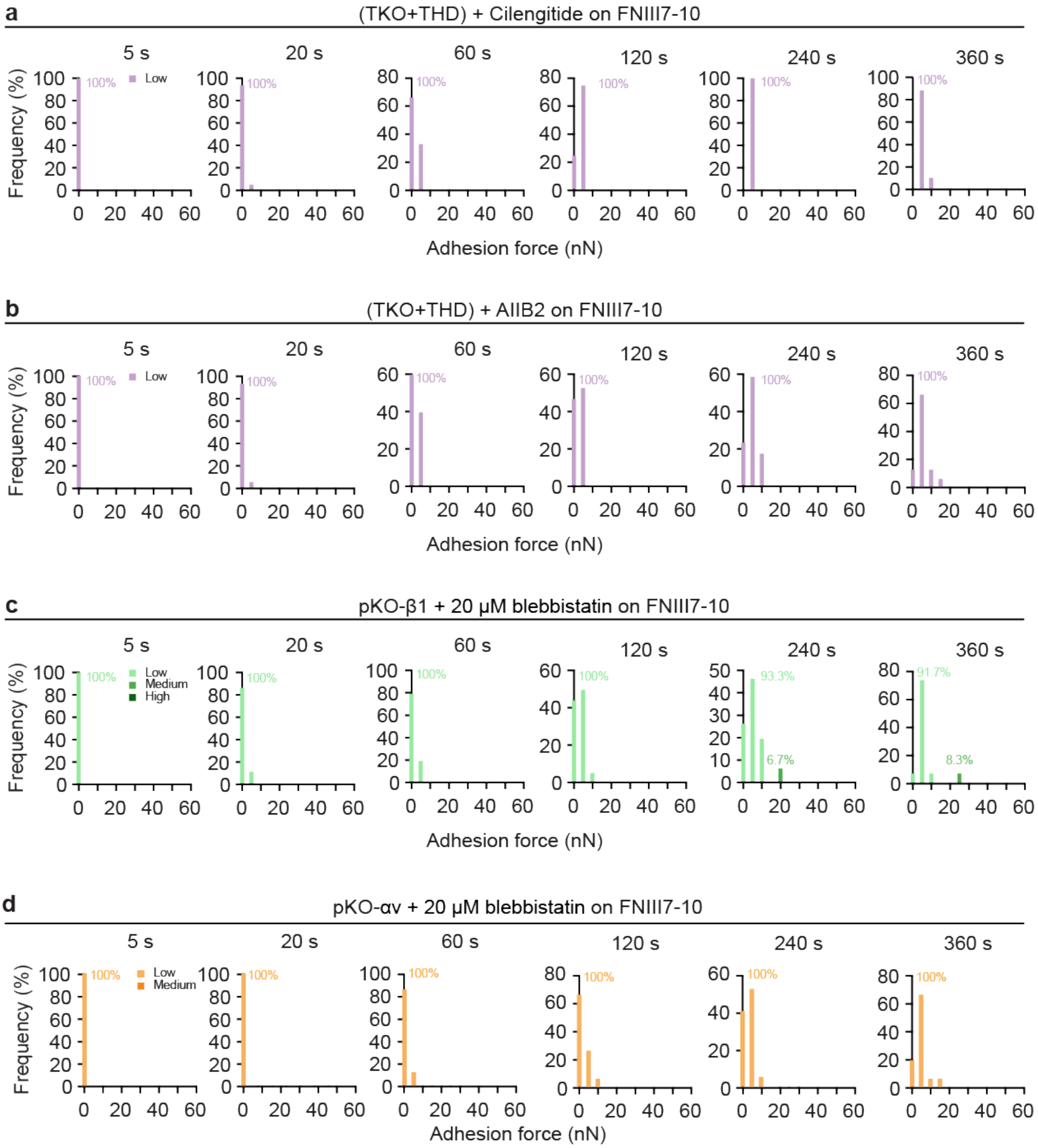


**Figure S3. Medium and high adhesion force populations depend on the engagement of integrins to the contractile actomyosin**. Adhesion force distribution of (a,b) talin1/2-depleted and talin1 head domain re-expressing (TKO+THD) fibroblasts in the presence of (a) cilengitide or (b) AIIB2 and (c,d) myosinII-contractility inhibited (c) pKO-β1 or (d) pKO-αv fibroblast. Histograms with a bin width of 5 nN show the adhesion force for the given contact time. The color of the bars indicate the adhesion force population as annotated. The numbers indicate the occupancy of the populations.

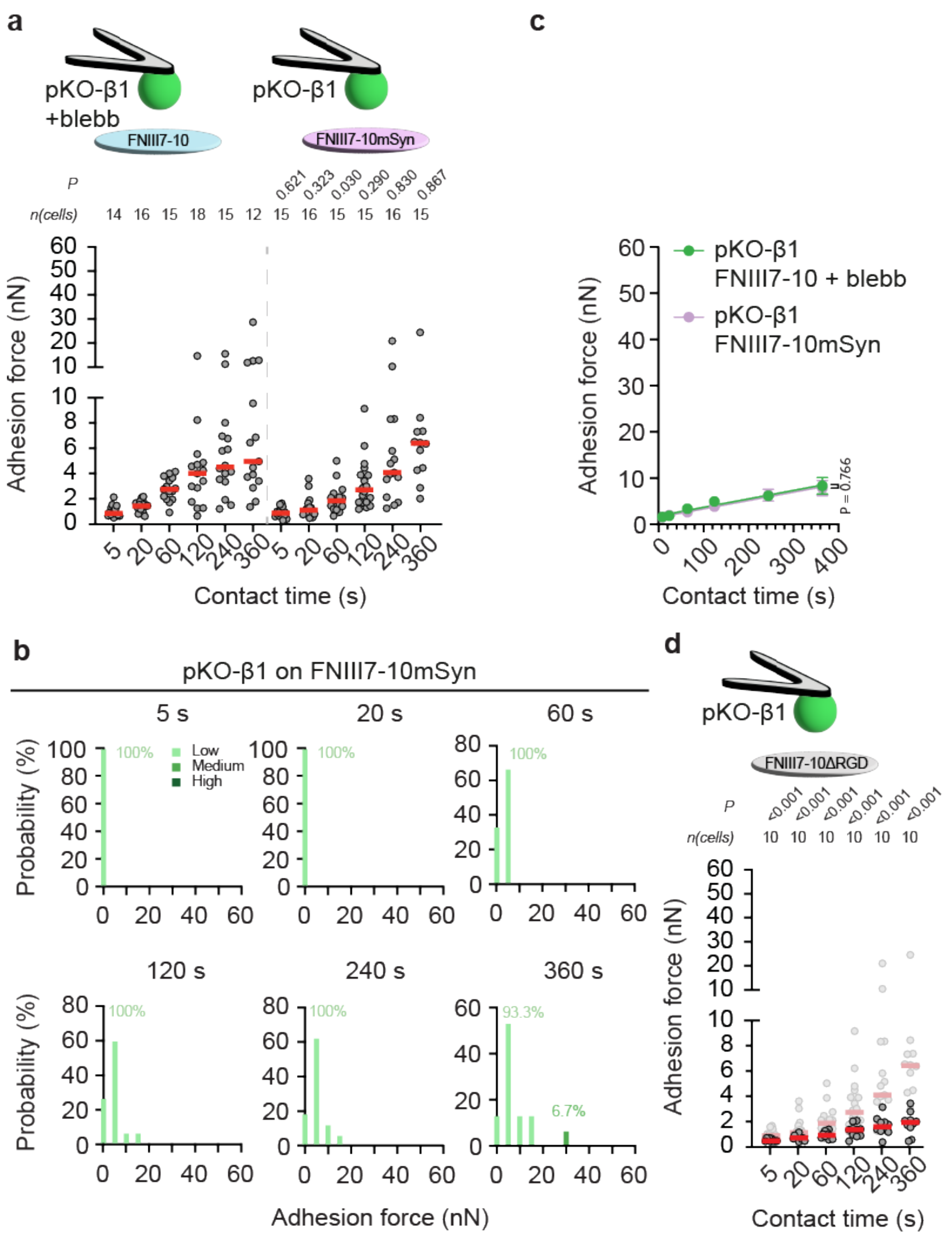


**Figure S4. The fast adhesion strengthening mode of pKO-β1 fibroblasts depends on the FN synergy site. a)** Adhesion forces of (left) 20 µM blebbistatin-treated pKO-β1 fibroblasts to FNIII7-10 (data taken from Fig. 2d) and (right) of untreated pKO-β1 fibroblasts to FNIII7-10mSyn (data taken from Fig. 3a). Dots represent adhesion forces of individual fibroblasts, red bars the median, and *n*(cells) the number of fibroblasts tested. *P* values comparing adhesion forces at the respective contact time were calculated using two-sided Mann-Whitney tests. **b)** Histograms with a bin width of 5 nN show the adhesion force distribution of pKO-β1 fibroblasts adhering to FNIII7-10mSyn for the given contact time. The color of the bars indicate the adhesion force population as annotated. The numbers indicate the occupancy of the populations. **c)** Adhesion strengthening of 20 µM blebbistatin-treated pKO-β1 fibroblasts to FNIII7-10 or pKO- β1 fibroblasts to FNIII7-10mSyn was quantified as a slope of a linear function (lines) fitted to all adhesion forces at all contact times. Dots represent the mean adhesion force and bars the standard error of the mean (SEM). The given *P* value compares the linear fit of both data sets and was calculated by extra sum-of-squares F-tests. **d)** Adhesion forces of pKO-β1 fibroblasts to FNIII7-10ΔRGD at given contact times. Adhesion forces of pKO-β1 fibroblasts to FNIII7-10mSyn (data taken from Fig. 3a) are given in grey as reference. Data representation as in a). P values comparing given and reference data were calculated using two-sided Mann-Whitney Tests.

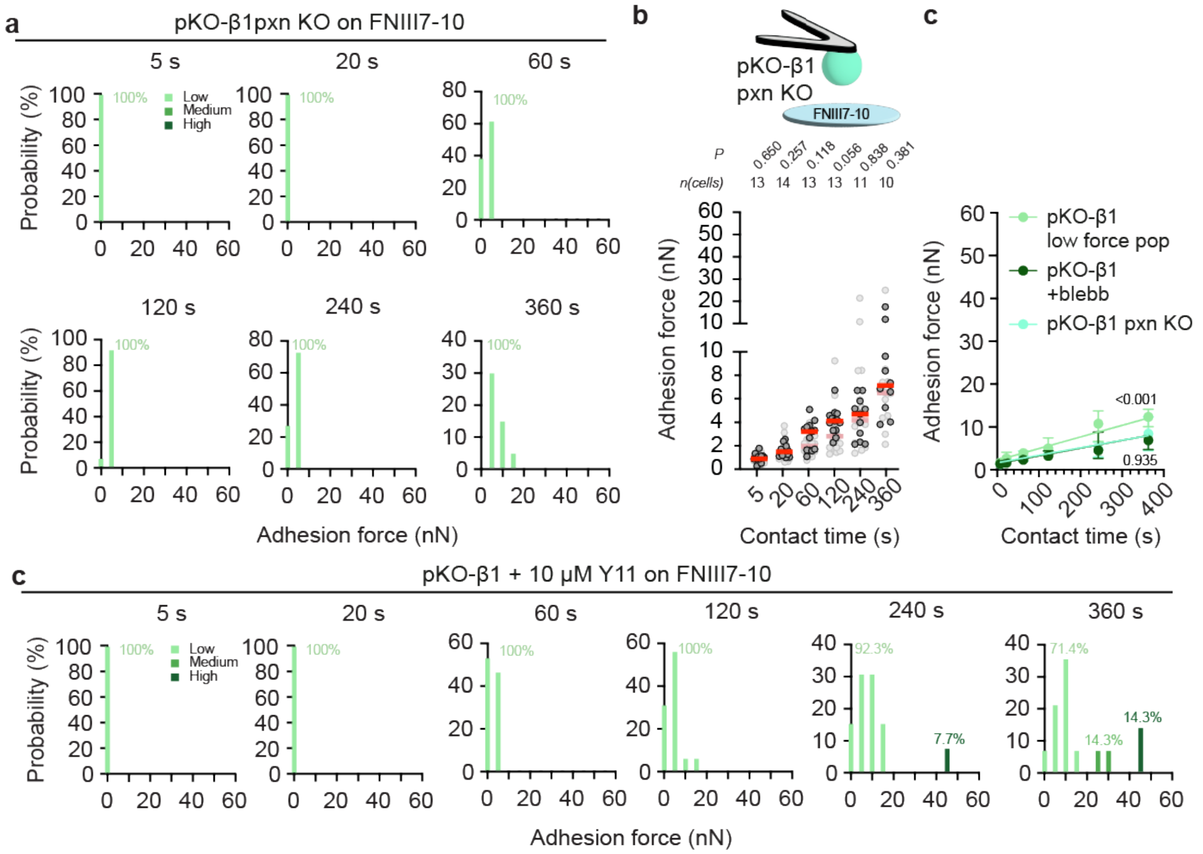


**Figure S5. Paxillin and FAK signaling are required for fast adhesion strengthening of pKO-β1 fibroblasts. a)** Histograms with a bin width of 5 nN show the adhesion force distribution of pKO-β1 pxn KO fibroblasts adhering to FNIII7-10 for the given contact time. Light green bars indicate the low adhesion force population, medium green bars the medium adhesion force population and dark green bars show the high adhesion force population. **b)** Adhesion forces of pKO-β1 pxn KO fibroblasts to FNIII7-10 (data taken from Fig. 5b). Adhesion forces of 20 µM blebbistatin-treated pKO-β1 fibroblasts adhering to FNIII7-10 (data taken from Fig. 4a) are given as grey as reference. Dots represent adhesion forces of individual fibroblasts, red bars the median, and *n*(cells) the number of fibroblasts tested. *P* values comparing adhesion forces at the respective contact time were calculated using two-sided Mann-Whitney tests. **c)** Adhesion strengthening of low adhesion force population pKO-β1 fibroblasts, 20 µM blebbistatin-treated pKO-β1 fibroblasts to FNIII7-10 or pKO-β1 pxn KO fibroblasts to FNIII7-10mSyn was quantified as a slope of a linear function (lines) fitted to all adhesion forces at all contact times. Dots represent the mean adhesion force and bars the standard error of the mean (SEM). The given *P* values compare the adhesion strengthening of (top) the low adhesion force population pKO-β1 fibroblasts to FNIII7-10 with of pKO-β1 pxn KO fibroblasts to FNIII7-10mSyn and (bottom) of the low adhesion force population pKO-β1 fibroblasts to FNIII7-10 and of blebbistatin-treated pKO-β1 fibroblasts to FNIII7-10. The P values were calculated by extra sum-of-squares F-tests. **d)** Histograms with a bin width of 5 nN show the adhesion force distribution of 10 µM Y11-treated pKO-β1 fibroblasts adhering to FNIII7-10 for the given contact time.

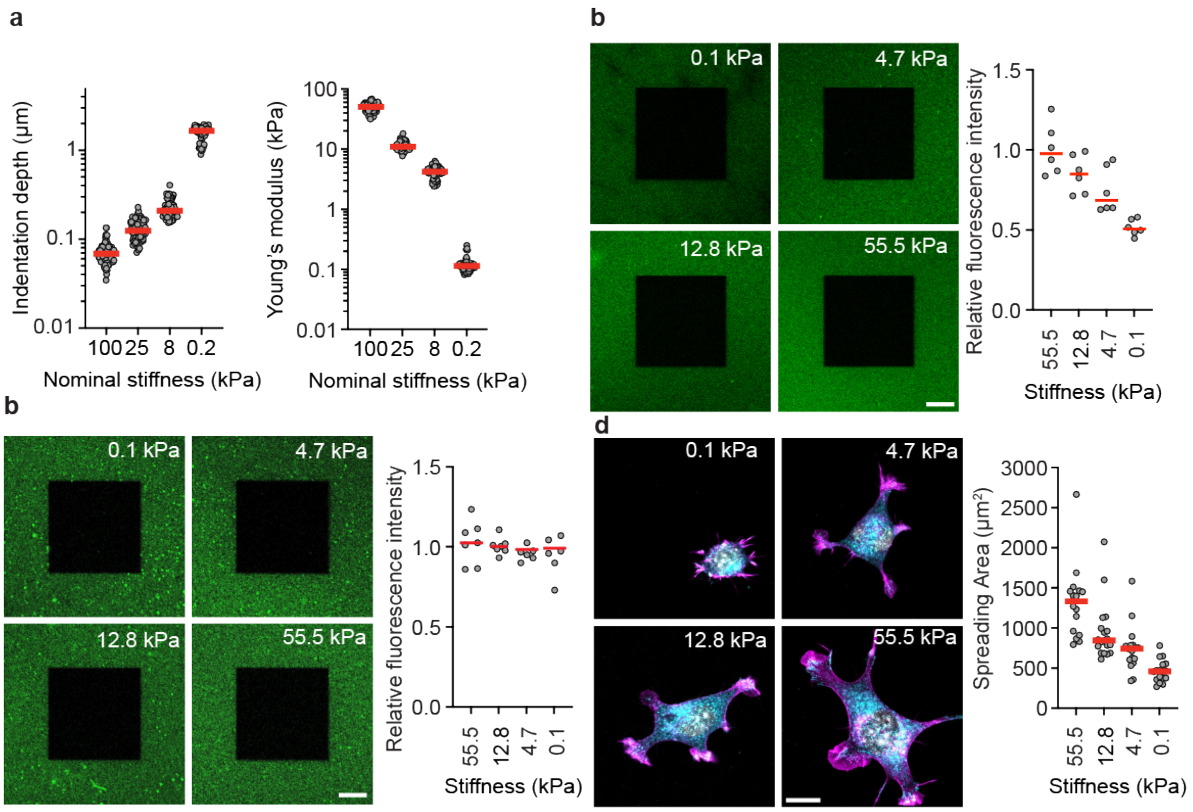


**Figure S6. Characterization of PAA gels for SCFS. a)** Indentation depth (left) and Young's moduli of PAA gels quantified with a 6 µm glass bead attached to a microcantilever to reach a force of 1 nN. Dots represent indentation depth or Young's modulus of each experiment and red bars the median. **b)** Left, representative immunofluorescent images of b) FN functionalized PAA gels or c) PAA gels functionalized with adjusted FN concentration having the given stiffness. The dark square in the middle of images shows a photobleached area to validate specific signal. Right, relative fluorescent intensity to the average intensity on the stiffest gel. Dots are the mean relative intensities of six line profiles across each images and red lines their median. **d)** Left, representative images of pKO-β1 fibroblasts spread for 90 min on FNIII7-10 functionalized PAA gels having the given nominal stiffness. Paxillin staining is shown in cyan, actin staining in magenta and the nucleus in white. Right, quantification of the spreading area of pKO-β1 fibroblasts by paxillin staining on differently stiff substrates. Dots represent spreading areas of individual fibroblasts and red bars the median. All scale bars, 20 µm.

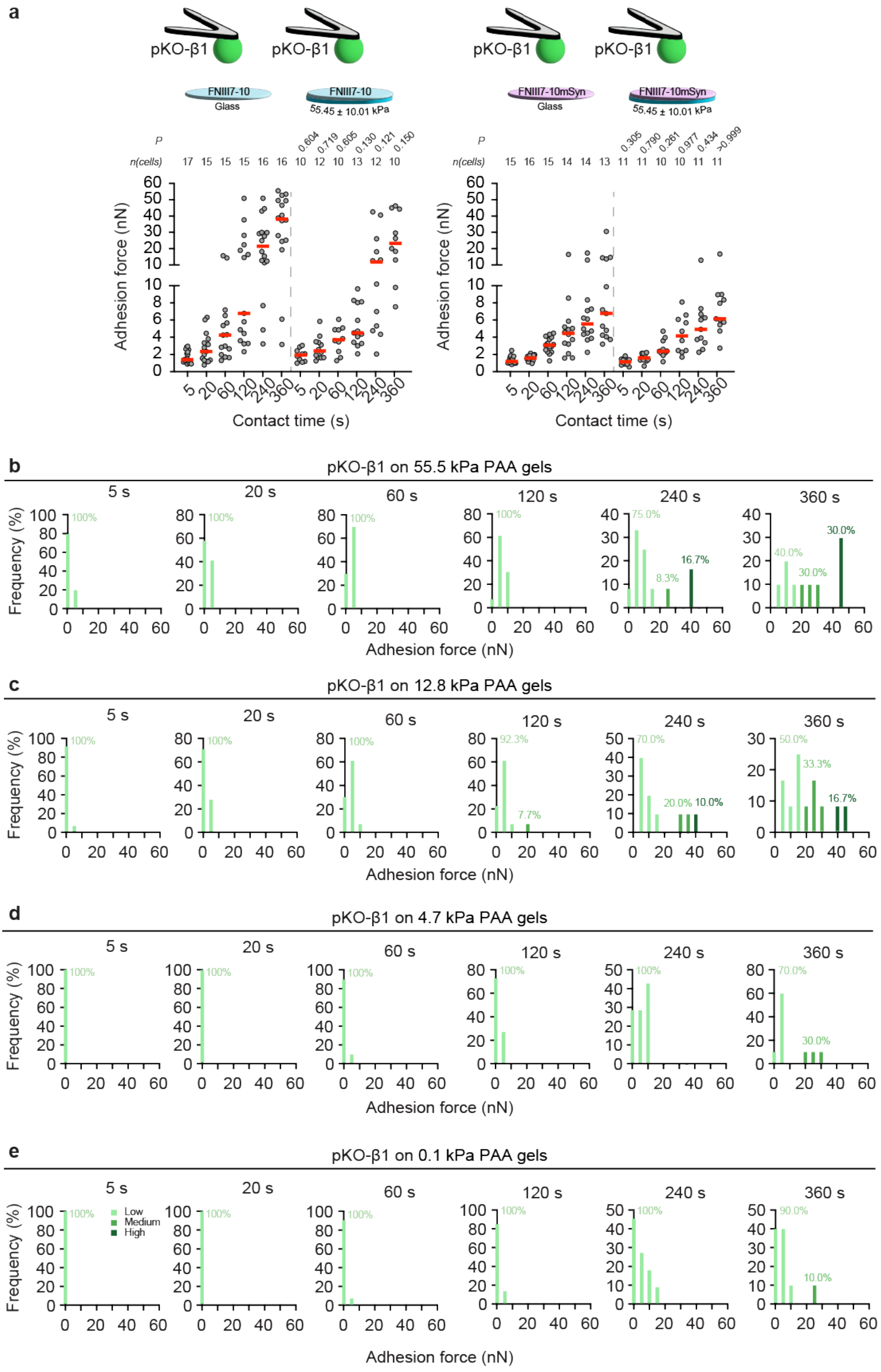
a
pKO-β1
FNIII7-10
Glass
55.45 ± 10.01 kPa
FNIII7-10mSyn
P
n(cells)
17 15 15 15 16 16 10 12 10 13 12 10
0.604 0.719 0.605 0.130 0.121 0.150
15 16 15 14 14 13 11 11 10 10 11 11
0.305 0.790 0.261 0.977 0.434 >0.999
Adhesion force (nN)
Contact time (s)
b
pKO-β1 on 55.5 kPa PAA gels
c
pKO-β1 on 12.8 kPa PAA gels
d
pKO-β1 on 4.7 kPa PAA gels
e
pKO-β1 on 0.1 kPa PAA gels
5 s
20 s
60 s
120 s
240 s
360 s
Frequency (%)
Adhesion force (nN)
Low
Medium
High

**Figure S7. α5β1 integrin populate the low, medium, and high adhesion force population substrate stiffness dependently.** a) Adhesion forces of pKO-β1 fibroblasts or adhering to (left) FNIII7-10-coated glass (data taken from Supplementary Fig. 2a) and FNIII7-10-functionalized 55.45 kPa PAA gels (data taken from Fig. 4a) or (right) FNIII7-10mSyn-coated glass (data taken from Fig. 3a) and FNIII7-10mSyn-functionalized 55.45 ± 10.01 kPa PAA gels (data taken from Fig. 4b). Dots represent adhesion forces of individual fibroblasts, red bar the median, and *n*(cells) the number of fibroblasts tested. *P* values comparing adhesion forces at the respective contact time were calculated using two-sided Mann-Whitney tests. **b-e)** Histograms with a bin width of 5 nN show the adhesion force distribution of pKO-β1 fibroblasts adhering to FNIII7-10-functionlaized PAA gels having given stiffness and for the given contact time. Light green bars indicate the low and medium dark green bars the medium adhesion force population. The numbers indicate the occupancy of the populations.
